# Hydraulic diversity stabilizes productivity in a large-scale subtropical tree biodiversity experiment

**DOI:** 10.1101/2021.01.06.425434

**Authors:** Florian Schnabel, Xiaojuan Liu, Matthias Kunz, Kathryn E. Barry, Franca J. Bongers, Helge Bruelheide, Andreas Fichtner, Werner Härdtle, Shan Li, Claas-Thido Pfaff, Bernhard Schmid, Julia A. Schwarz, Zhiyao Tang, Bo Yang, Jürgen Bauhus, Goddert von Oheimb, Keping Ma, Christian Wirth

## Abstract

Extreme climatic events threaten forests and their climate mitigation potential globally. Understanding the drivers promoting ecosystem stability is therefore considered crucial to mitigate adverse climate change effects on forests. Here, we use structural equation models to explain how tree species richness, asynchronous species dynamics and diversity in hydraulic traits affect the stability of forest productivity along an experimentally manipulated biodiversity gradient ranging from 1 to 24 tree species. Tree species richness improved stability by increasing species asynchrony. That is, at higher species richness, inter-annual variation in productivity among tree species buffered the community against stress-related productivity declines. This effect was mediated by the diversity of species’ hydraulic traits regarding drought tolerance and stomatal control, but not by the community-weighted means of these traits. The identified mechanisms by which tree species richness stabilizes forest productivity emphasize the importance of hydraulically diverse, mixed-species forests to adapt to climate change.

## Introduction

Climate change is increasing the frequency and severity of droughts and other extreme events, threatening tree growth and survival globally^1^, including in comparably humid tropical and subtropical forests^2^. This compromises the ability of the world’s forests to act as a carbon sink^3^ and as a nature-based solution to climate change^4^. Stability, the ability of forests to maintain functioning in periods of stress, is consequently emerging as a primary focus of forest management in the 21st century. One key management strategy to enhance stability may be to increase tree species richness in secondary and plantation forests^5–7^. However, we lack a comprehensive understanding of what drives biodiversity–stability relationships in forest ecosystems.

There is compelling evidence that species richness can stabilize community biomass production against variable climate conditions such as droughts or extremely wet years^8–11^. However, the majority of this evidence comes from grassland ecosystems. Biodiversity–stability relationships likely differ between forests and grasslands because trees invest in long-lasting structures and therefore community composition changes more slowly in forests^7^. The few existing studies in forests support the notion that species richness stabilizes aboveground wood production, hereafter referred to as ‘productivity’, of mixed-species tree communities^7,12–14^, but the underlying mechanisms remain largely unknown.

According to the insurance hypothesis^15,16^ a mixture of tree species with different strategies should help to maintain or increase the functioning of forests under highly variable climatic conditions, thus increasing their temporal stability. This stability^17^ is often quantified as temporal mean productivity (µ) divided by the temporal standard deviation in productivity (σ)^e.g.8,9^ and may be promoted by diversity in mixed-species tree communities via two principal mechanisms^15^. First, overyielding, which refers to an increased temporal mean productivity in mixtures compared to monocultures, has been reported by numerous studies in natural and experimental forests^18–21^. Here, different species perform relatively better in mixtures than in monocultures, for example through complementary resource use or facilitation and this higher performance can increase stability^7^. Second, decreased temporal variation in community productivity through buffering of the effects of stress may increase stability. In contrast to overyielding, little is known about this buffering effect of biodiversity in forest ecosystems. Various mechanisms may decrease temporal variation in productivity^15,17,22,23^ but arguably the one most supported by theoretical and observational studies in grasslands and increasingly also in forests is species asynchrony^8,22,24^. In forests, these asynchronous inter-annual dynamics in productivity among tree species (hereafter ‘species asynchrony’^23^) have been found to be the strongest driver of community stability^7,12–14,25^. Asynchronous species dynamics may result from intrinsic rhythms like phenology or mast seeding^26,27^, differential responses of species to extrinsic factors such as climatic conditions^23,28^ and species interactions in mixtures like resource partitioning or biotic feedbacks^29,30^. Species asynchrony may buffer the temporal variation in community productivity during times of stress as some species likely maintain functioning or compensate for the productivity losses of other species (Fig. 1). This stabilizing effect may be especially important in the context of the global increase in the severity and frequency of drought events^31,32^. Hence, there is an urgent need to identify the characteristics that allow tree species and species mixtures to maintain functioning under future drought conditions.

**Fig 1.**
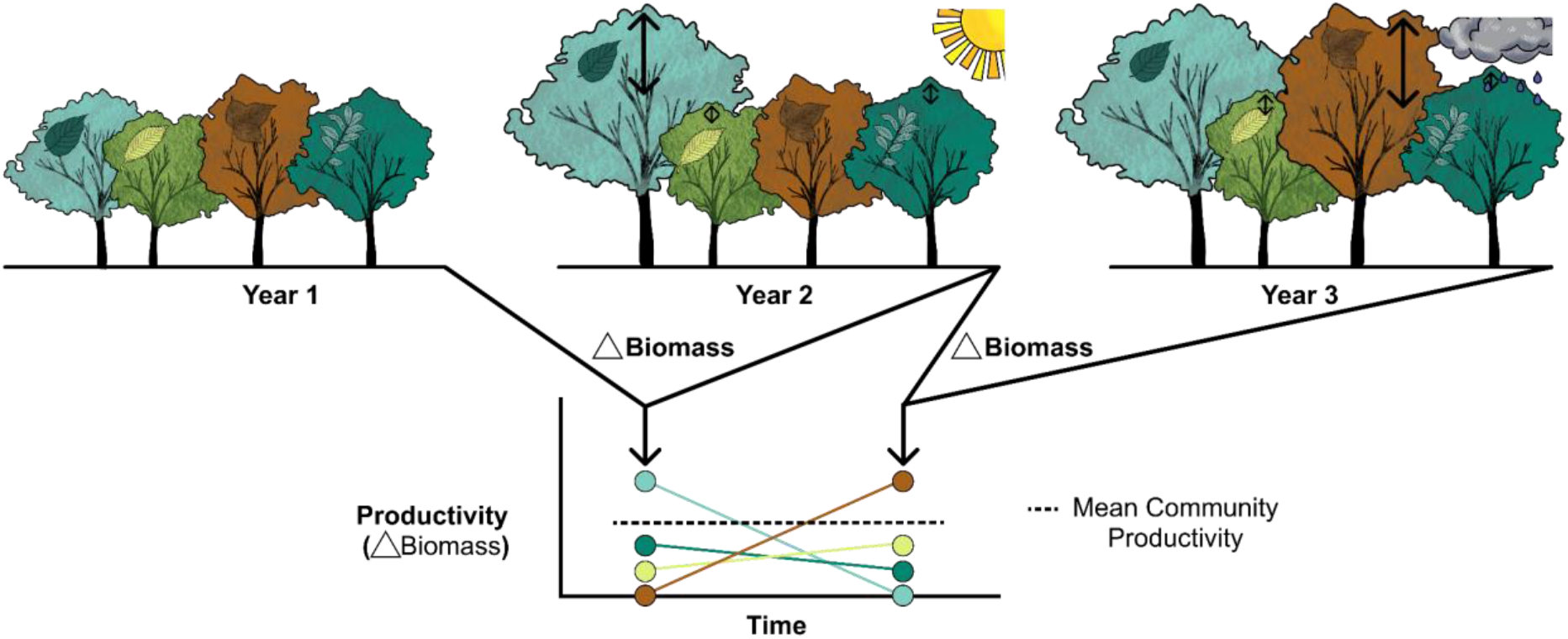
Graphical illustration of asynchronous species responses in mixed-species tree communities to contrasting climatic conditions over a period of three years. The tree community experiences a ‘normal’ (year 1), an exceptionally dry (year 2) and an exceptionally wet (year 3) year, which result in distinctly different growth responses of the participating species but the same community productivity due to compensatory dynamics. In our hypothetical example taken from a four species mixture in the BEF-China experiment, one species (*Nyssa sinensis*, light turquoise) does not close its stomata fast during water shortage (water-spender) and might grow well during drought, a second species (*Liquidambar formosana*, brown) exhibits a fast downregulation of stomatal conductance at increasing water pressure deficits and its productivity is thus more strongly reduced during drought (water-saver), while the two other species (*Castanea henryi, Sapindus mukorossi*) do not show strong reactions to the changing climatic conditions. The reverse response pattern is found during an exceptionally wet year. We hypothesize here that such asynchronous species dynamics are the key driver behind stabilizing effects of species richness on productivity in mixed-species forests and that the functional traits of co-existing species — especially those associated with hydraulic functioning — may help to elucidate the mechanisms that produce this species asynchrony.

While the number of species may increase stability, communities also require certain hydraulic characteristics to endure drought. Among other features such as non-structural carbohydrates^33^, two key hydraulic strategies that determine a tree’s response to drought are drought tolerance and stomatal control^28^. First, drought tolerance depends on xylem resistance to cavitation because embolism decreases water availability and may ultimately lead to desiccation and tree death^2,28^. Here, we use the threshold at which 50% of xylem conductivity is lost due to cavitation (Ψ_50_; measured as water potential) as key trait^2^ and, in addition, classic traits of the leaf economics spectrum (indicating conservative vs acquisitive resource use)^34^ to quantify drought tolerance. Second, tree species may follow different strategies of stomatal control. Some rely on continued water extraction and keep their stomata open, i.e. they continue to transpire even though this poses a high risk for cavitation-induced mortality under extreme drought (called water-spenders or anisohydric species)^28,35,36^. Other tree species decrease their stomatal conductance quickly during water shortage to avoid transpiration losses and xylem cavitation but may risk carbon starvation under prolonged droughts (called water-savers or isohydric species). Consistent with recent perspectives^36^, we view stomatal control here along a gradient from water-spending to water-saving species behaviour and quantify it through physiological traits such as stomatal conductance and control of conductance under increasing water pressure deficits^37,38^.

These different hydraulic strategies may enable mixed-species forests to stabilize community productivity in two ways. First, tree species richness may increase stability indirectly via promoting species asynchrony through diversity in functional traits related to hydraulic water transport (hereafter ‘hydraulic diversity’^39^). The importance of tree species richness and species asynchrony for stability is supported by previous studies^7,12–14^. However, these studies were based on observational data from naturally assembled forests (with only one exception^14^) and tree species richness gradients were short. Therefore, it remains difficult to establish causal relationships between tree diversity and stability. In particular, the mechanistic links between tree species richness, species asynchrony and stability as well as the underlying trait-based mechanisms remain unknown for forests. Second, stability could also be influenced by the community-weighted mean of hydraulic traits, as indicated by findings in grassland diversity experiments where stability was higher in communities dominated by species with traits associated with conservative resource use^8^. However, as we expect species asynchrony to be the key driver of stability in forests, we consider trait diversity more important for stability than community trait means.

We use structural equation models (SEMs) to test the direct and indirect effects of species richness, species asynchrony, hydraulic diversity and community-weighted means of hydraulic traits on the stability of community productivity under the controlled conditions of a large-scale tree biodiversity experiment (BEF-China^18,40^; biodiversity–ecosystem functioning China). Our experiment is located in the highly diverse subtropical forests of China and features a gradient of species richness ranging from monocultures up to mixtures of 24 tree species planted at two sites using multiple species pools. All species occurred at all richness levels, thus avoiding any confounding between species occurrence and richness. In our study, stomatal control and drought tolerance strategies form two orthogonal species trait gradients (Supplementary Fig. 1), which allows us to quantify their relative contributions to species asynchrony and stability. Specifically, we tested the following hypotheses:

**H1** Tree species richness increases stability via species asynchrony.

**H2** Hydraulic diversity in stomatal control and drought tolerance strategies increases stability through species asynchrony.

**H3** Stability increases through buffered temporal variation in productivity mediated by species asynchrony and by overyielding.

## Results

Overall, the stability of community productivity significantly increased with species richness in our experimental tree communities. We found significant positive relationships between stability, species asynchrony and hydraulic diversity – calculated as functional dispersion of species along two trait gradients related to stomatal control (functional diversity of stomatal control) and resistance to cavitation (functional diversity of drought tolerance) (Fig. 2; Supplementary Figs. 1-4; Supplementary Table 1–2). In contrast, community-weighted means (CWMs) of these hydraulic gradients did not influence the stability of productivity (Supplementary Fig. 5; Supplementary Table 2). Specifically, we found a significant positive effect of species richness on stability (t=3.98, P<0.001, n=375; Fig. 2), which was insensitive to the inclusion or exclusion of monocultures into the models (Supplementary Fig. 6; Supplementary Table 2). Among the analysed bivariate relationships species asynchrony had, as predicted, the strongest positive effect on stability in mixtures and explained most of its variation (t=10.13, P<0.001, marginal R^2^=34%, n=218; Fig. 2; Supplementary Table 2). Species asynchrony significantly increased with species richness (t=9.53, P<0.001), functional diversity of stomatal control (t=5.29, P<0.001) and functional diversity of drought tolerance (t=5.84, P<0.001) (Supplementary Figs. 2-3). Direct effects of hydraulic diversity on stability were weak: we found a marginally significant positive effect of functional diversity of stomatal control on stability (t=1.92, P=0.058) but no significant relationship with functional diversity of drought tolerance (t=1.12, P=0.27; Supplementary Fig. 4). Hydraulic diversity explained a much higher share of variability in asynchrony than it did in stability (Supplementary Table 2).

**Fig 2.**
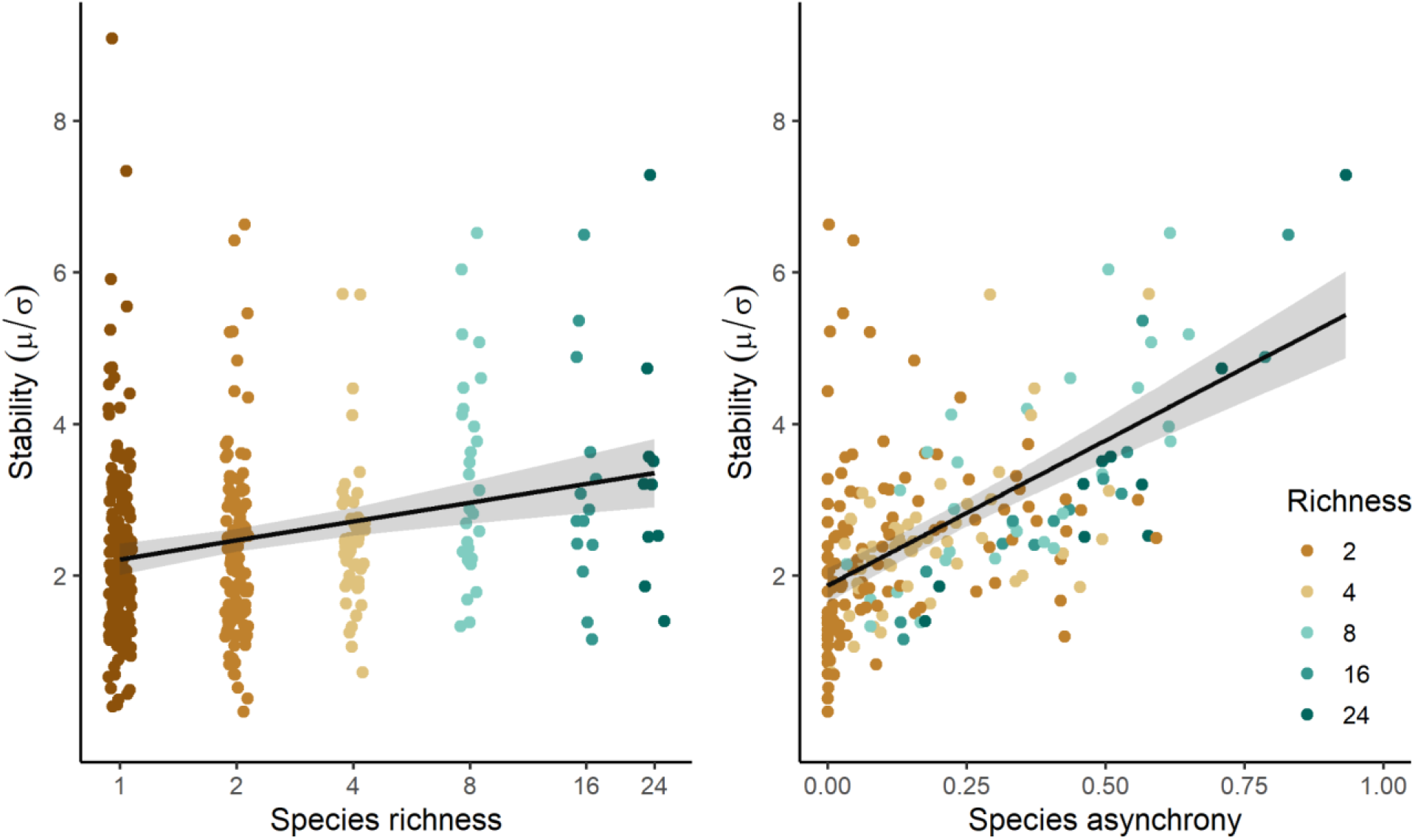
Effects of species richness and species asynchrony on stability. Lines are linear mixed-effect model fits that show (a) significant increases in stability with species richness (P<0.001) along a planted diversity gradient ranging from monocultures up to mixtures of 24 tree species and (b) significant increases in stability with species asynchrony (P<0.001) in mixtures. Species asynchrony ranges from 0 to 1, where 0 represents complete synchrony and 1 complete asynchrony. Grey bands represent a 95% confidence interval. See Supplementary Table 2 for details on the fitted models.

Structural equation models allowed us to disentangle the direct and indirect drivers and connections behind observed diversity effects on stability (Fig. 3). Species asynchrony was the principal mediator of indirect effects of species richness via hydraulic diversity on stability. Our model fit the data well (Fishers’ C=9.7, d.f.=8, P=0.28, n=218). The hypothesized pathways explained 35% of variation in stability (fixed effects, marginal R^2^), which increased to 58% if both fixed and random effects (conditional R^2^) were considered. Species richness, functional diversity of stomatal control and functional diversity of drought tolerance explained 52% of variation in species asynchrony (marginal R^2^). Species asynchrony was the strongest direct driver of stability (standardized path coefficient of direct effect 0.76, P<0.001). Tree species richness increased stability indirectly through increasing species asynchrony (standardized path coefficient of compound effect 0.35, i.e. the product of the coefficients along the path). We did not find an additional independent effect of species richness on stability (P=0.31), indicating that species asynchrony was the principal mediator of species richness effects on stability. Quantifying hydraulic diversity allowed us to disentangle some of the functional drivers behind asynchronous species responses: both functional diversity of stomatal control and functional diversity of drought tolerance contributed to increased stability via positive effects on species asynchrony (standardized path coefficients of direct effects on asynchrony 0.18, P=0.005 and 0.30, P<0.001, respectively). Functional diversity of drought tolerance also had a direct negative effect (P=0.007) on stability, that was smaller than its positive effect on asynchrony (standardized path coefficients of direct effects −0.21 vs 0.30). Importantly, only functional diversity but not community-weighted means of the hydraulic trait gradients explained variations in stability (effect of CWM of stomatal control and CWM of drought tolerance on stability both not significant with P≥0.25; Fig. 3).

**Fig 3.**
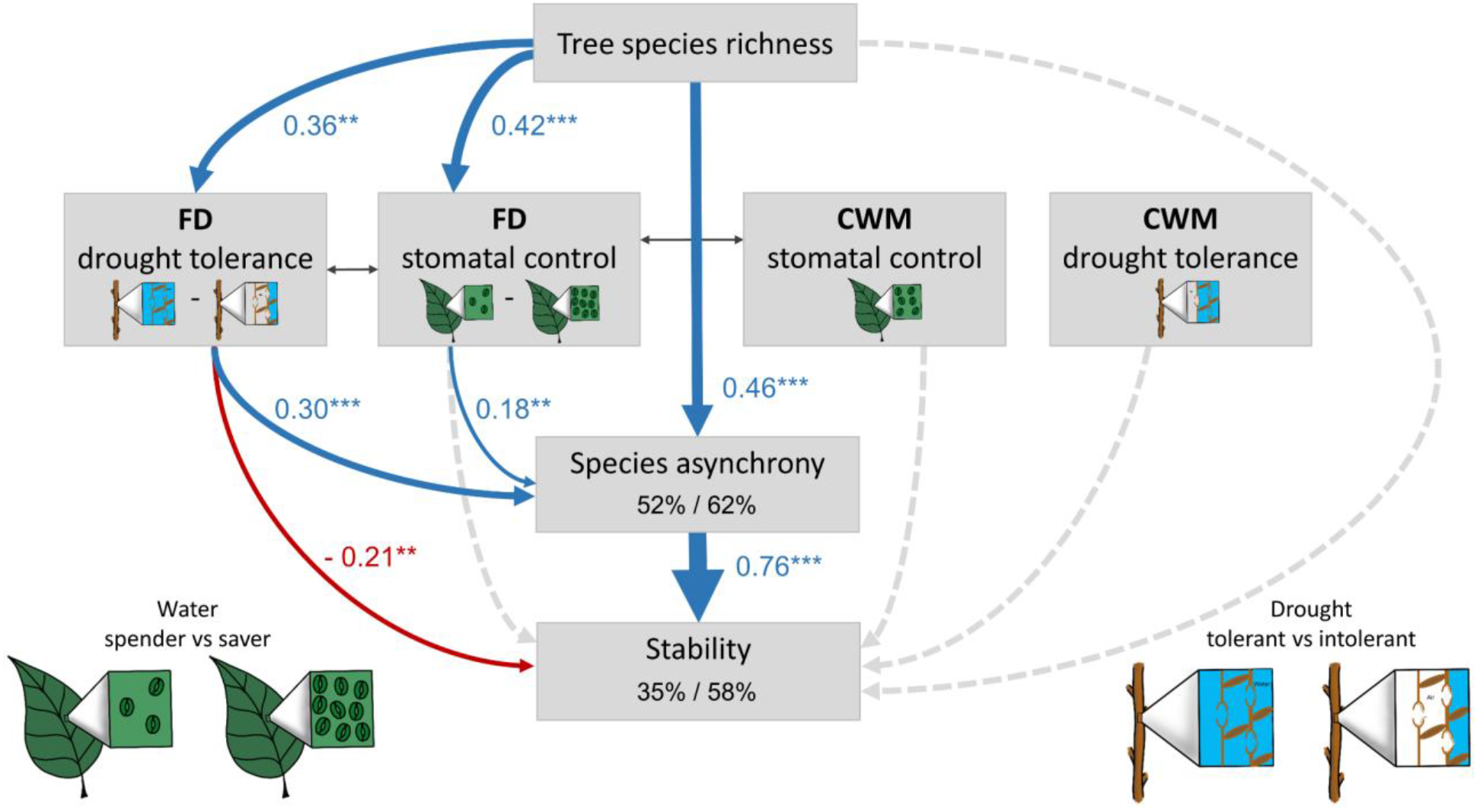
Direct and indirect effects of species richness, hydraulic diversity and community hydraulic means on stability. The structural equation model (SEM) tests the direct effects of tree species richness, functional diversity of stomatal control (FD stomatal control) and functional diversity of drought tolerance (FD drought tolerance) as well as their indirect effects mediated via species asynchrony on stability. Effects of community-weighted mean (CWM) traits are explored through testing the effect of the CWM of stomatal control (CWM stomatal control) and the CWM of drought tolerance (CWM drought tolerance) on stability. The sketches schematically illustrate the trait gradients: water-spending vs water-saving stomatal control (few vs abundant stomata) and drought tolerance (high vs low cavitation resistance). Functional diversity was calculated as abundance-weighted functional dispersion. The SEM fit the data well (Fisher’s C=9.7, P =0.28, d.f.=8, n=218 plots). Data is based on a long, experimental species richness gradient with mixtures of 2, 4, 8, 16 and 24 tree species. Examined variables are shown as boxes and relationships as directional arrows with significant positive effects in blue, significant negative effects in red and non-significant paths in dotted grey based on a hypothesis driven SEM framework (Supplementary Fig. 7). Standardized (significant) path-coefficients are shown next to each path with asterisks indicating significance (* P<0.05, ** P<0.01, *** P<0.001), path-width is scaled by coefficient size. Significant partial correlations^41^ are shown through grey, bi-directional arrows. The variation in species asynchrony and stability explained by fixed (left, marginal R^2^) and fixed together with random model effects (right, conditional R^2^) is shown in the corresponding boxes.

We further separated the components of our stability measure — the temporal mean (µ_AWP_) and the temporal standard deviation (σ_AWP_) of productivity — to examine the underlying cause of the observed biodiversity–stability relationships (Fig. 4). Tree species richness directly increased both the mean and the standard deviation of productivity similarly (standardized path coefficients of direct effects 0.23 and 0.30, respectively). Tree species richness thus increased mean productivity but this was accompanied by increased variation in productivity. However, species richness also decreased the standard deviation of productivity indirectly via its positive effect on species asynchrony with about the same strength (indirect effect of species richness on σ_AWP_ −0.3, calculated as the product of the coefficients along each significant path and their sum^41^; Fig. 4). Species asynchrony, which increased with species richness and hydraulic diversity, hence stabilized productivity through buffering its temporal variation (standardized path coefficient of direct effect of species asynchrony on σ_AWP_ −0.47, P<0.001). The unexpected direct negative effect of functional diversity of drought tolerance on stability (Fig. 3, see above) can be attributed to its positive effect on the temporal standard deviation (Fig. 4, marginally significant, P=0.06). Finally, the CWM of drought tolerance increased both, mean productivity and the standard deviation of productivity (standardized path coefficients of direct effects 0.21 and 0.16, respectively). That is, communities dominated by drought-intolerant species (those with higher trait gradient scores; Supplementary Fig. 1) had a higher productivity but tended to also have a higher variation in productivity. Overall, stability increased with species richness (Fig. 3) through increased mean productivity (i.e. overyielding) and buffered temporal variation in productivity (Fig. 4).

**Fig 4.**
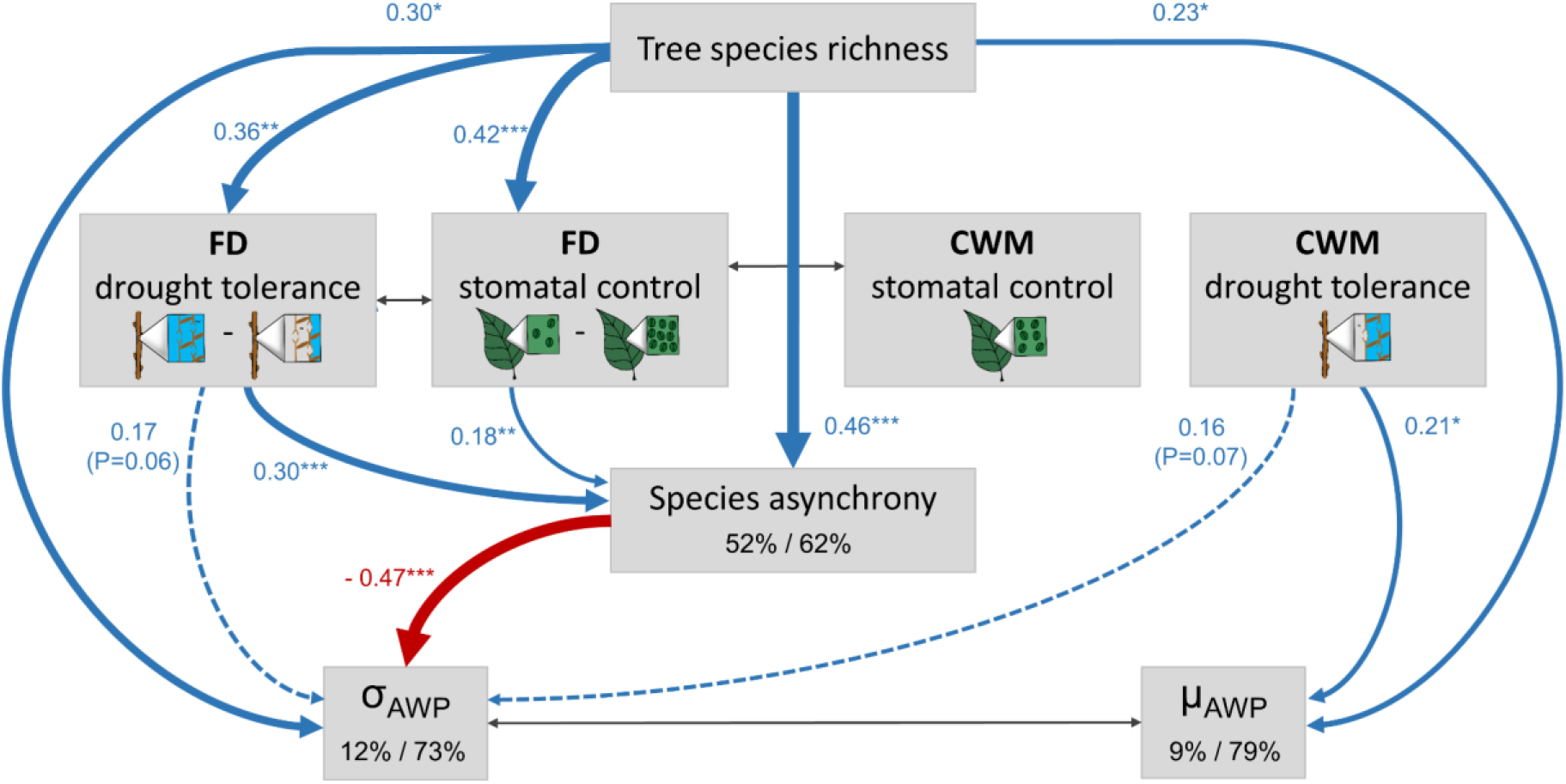
Structural equation model (SEM) of direct and indirect effects of species richness, species asynchrony, hydraulic diversity and community hydraulic means on the two components of stability, the temporal mean (µ_AWP_) and the temporal standard deviation of productivity (σ_AWP_), which represent overyielding and variance buffering effects, respectively. Increases in µ_AWP_ enhance stability through overyielding — a higher productivity in mixtures vs monocultures — and decreases in σ_AWP_ enhance stability through buffered variations in productivity. All drivers hypothesized to influence stability, i.e. species richness, functional diversity of stomatal control (FD stomatal control), functional diversity of drought tolerance (FD drought tolerance), CWM of stomatal control (CWM stomatal control), CWM of drought tolerance (CWM drought tolerance) and species asynchrony, were tested for their effects on µ_AWP_ and σ_AWP_. Only significant pathways (P<0.05) are shown here to avoid overplotting (see Supplementary Fig. 8 for the full model). The sketches schematically illustrate the trait gradients: water-spending vs water-saving stomatal control (few versus abundant stomata) and drought tolerance (high versus low cavitation resistance). The SEM fit the data well (Fisher’s C=9.7, global P=0.28, d.f.=8, n=218 plots). Data is based on a long, experimental species richness gradient with mixtures of 2, 4, 8, 16 and 24 tree species. Examined variables are shown as boxes and relationships as directional arrows with significant positive effects in blue, significant negative effects in red and non-significant paths in dotted grey. Standardized (significant) path-coefficients are shown next to each path with asteri sks indicating significance (* P<0.05, ** P<0.01, *** P<0.001), path-width is scaled by coefficient size. Significant partial correlations^41^ are shown through grey, bi-directional arrows. The variation in species asynchrony, µ_AWP_ and σ_AWP_ explained by fixed (left, marginal R^2^) and fixed together with random model effects (right, conditional R^2^) is shown in the corresponding boxes.

## Discussion

Our results provide experimental evidence that the insurance effect^15^ of diversity stabilizes tree productivity in forest ecosystems. We show that the stability of forest community productivity increases with tree species richness and that asynchronous productivity of co-existing species in response to climatic variation is the principal mediator of this diversity effect. As hypothesized, both hydraulic diversity in stomatal control and drought tolerance had net positive, indirect effects on stability that operated via enhanced species asynchrony. In contrast, the community-weighted means of these hydraulic trait gradients did not influence stability.

### Species asynchrony and stability

The controlled diversity gradient of the BEF-China experiment^40^ ranging from monocultures to mixtures of 24 tree species, detailed trait information and the use of structural equation models^41^ allowed us to disentangle the direct and indirect drivers of stability in forests in the absence of confounding environmental variation typically hampering interpretations in observational studies. We show here that species richness increases stability indirectly via promoting asynchronous species productivity over time. Stability and species asynchrony were positively correlated with tree species richness in former studies^7,12–14^. Our experimental results add support for the hypothesized causality in these studies and demonstrate that species richness can drive species asynchrony and thereby stability in highly diverse subtropical forests. Asynchronous productivity integrates different mechanisms, such as those captured by the selected hydraulic traits that help species to cope in different ways with the variable climatic conditions typical for the sites (Supplementary Fig. 9). This species asynchrony due to diverse hydraulic strategies enhanced stability via buffering (reducing) variation of productivity over the 10-year observation period^7,8,13,15,23^. Species richness also increased temporal mean productivity directly. This finding is in line with a rapidly increasing number of studies reporting that forest productivity increases with increasing tree species richness^14,18–20^. However, this increased productivity by itself did not increase stability because species richness also increased the temporal variation of productivity. Stability only increased due to the variance buffering effect of species asynchrony on productivity (Fig. 4).

The asynchronous growth dynamics of different species in our experimental tree communities likely result from different, non-mutually exclusive mechanisms. First, extrinsic factors like climate may increase species asynchrony. Species react differently to climatic conditions^e.g.2,28^ and species asynchrony is thus likely driven by differential growth responses of species in mixture to climatic variability. Next, tree growth in mixtures is shaped by tree–tree interactions such as resource partitioning and biotic feedbacks^21,29,30,42^ which may in turn be modulated by variation in climatic conditions^43–45^. Finally, intrinsic rhythms like mast seeding can influence species productivity and its inter-annual variability^26^ thereby inducing species asynchrony. These intrinsic factors are, however, presumably less important in young forest stands. We thus expect that the observed strong species asynchrony resulted from differential response strategies of species to inter-annual variation in climatic conditions (the only environmental variable with strong inter - annual variation during our 10-year study period; Supplementary Fig. 9) and how these strategies, which we quantified via hydraulic traits, shape the nature of tree–tree interactions between years with different climatic conditions.

### Hydraulic diversity and stability

We used two orthogonal hydraulic trait gradients (Supplementary Fig. 1), related to species - specific stomatal control and drought tolerance and explored their relative contribution to stability. This allowed us to explain some of the mechanisms that induced asynchronous growth dynamics and stabilized productivity in the face of highly variable climatic conditions. According to a contextualization of the stress-gradient hypothesis’ for forests^46^, complementary species interactions likely increase in frequency and intensity with decreasing resource availability. Hence, while we show that species asynchrony increased with dissimilarity in hydraulic traits, we expect that the relative importance of these traits for tree productivity is higher in dry years than in years with ample water supply. Consistent with this expectation, we found the strongest positive tree species richness effects on tree growth during severely dry years in former studies^14,43^. This climate-driven biodiversity effect was modulated by a species drought tolerance at our study site (quantified as in this study as resistance to cavitation; Ψ_50_)^43^. Hence, we consider the hydraulic traits used here to be suitable response traits that capture inter-annual changes in productivity as driven by inter-annual variation in climatic conditions. This is in line with the ubiquity of vulnerability to drought across all forest ecosystems^2^, including comparably humid subtropical forests.

Functional diversity in stomatal control increased species asynchrony and thus indirectly stability through reducing variation in productivity. This effect of hydraulic diversity on stability is consistent with recent evidence that tree hydraulic diversity buffers temporal variations in forest ecosystem carbon fluxes during drought^39^. Functional diversity in stomatal control may promote species asynchrony among water spenders and water savers. The former keep their stomata open and continue to transpire during drought. This strategy, however, likely relies on continuous water uptake via roots to balance transpiration losses and carries high cavitation risks^28,36^, a principle mechanism behind drought-induced mortality across tree taxa^47^. Conversely, water savers can reduce this risk but may face carbon starvation under prolonged droughts^28^ even though starvation is less ubiquitous than cavitation^47^. These contrasting stomatal control strategies themselves may induce strong inter-annual changes in tree growth while also determining the water availability in mixed-stands through soil water partitioning between co-existing species^48,49^. In tree neighbourhoods comprising species with different stomatal control strategies, water spenders may benefit from soil water left by their water-saving neighbours during drought, while water savers may capitalize on improved soil water conditions after a drought due to their potentially faster drought recovery^48^. Both may thus profit from neighbourhood species richness as the likelihood for functional dissimilar species increases with species richness. Diversity in stomatal control is therefore one potential mechanism that explains reported positive neighbourhood species richness effects on individual tree growth especially during drought^14,43^ but conceivably also during wetter years.

Functional diversity in drought tolerance had a stronger positive influence on stability via asynchronous species productivity but also a direct (albeit weaker) negative effect on stability. Whereas drought-tolerant species can stabilize productivity of mixed-species communities through lower risks for xylem cavitation during dry years^2,28^, the latter, characterized by traits associated with an acquisitive resource use strategy^37^ (see Supplementary Fig. 1), can stabilize productivity in wet years. This acquisitive resource use may, moreover, enable soil water partitioning between neighbours during dry years in favour of drought-intolerant species^43^. The reasons for the additional direct negative effect of drought tolerance diversity on stability remain speculative. Nevertheless, as it resulted from increased temporal variation in community productivity at higher species richness (Fig. 4), it might be related to a dieback of highly drought - sensitive species in drought years destabilizing community productivity.

The direct positive effects of species richness on species asynchrony, remaining after accounting for the indirect effects via hydraulic diversity, may result from dissimilarity in traits^8^ that were not considered here such as leaf phenology^27^, storage of non-structural carbohydrates^33^, traits regulating biotic feedbacks^29^ and below- and aboveground structural traits^50–52^. For example, rooting-depth, complementary water uptake through niche differentiation^53^ and facilitation via hydraulic redistribution^48^ between species could be important drivers of species asynchrony and stability belowground.

### Community hydraulic means and stability

In contrast to hydraulic diversity, we did not find effects of community-weighted means of hydraulic traits on stability. Species asynchrony, the key driver of stability, depends naturally more on diverse species strategies (see Fig. 1) than on the prevalence of a specific strategy within a community. The absence of community mean effects on stability and the preponderance of negative selection effects developing over time in our experiment^18^ underlines that the observed responses are not simply related to communities becoming increasingly dominated by particularly stable-growing species with stand development. We found some indication for increased productivity in communities dominated by rather drought-intolerant (acquisitive) species, consistent with the common expectation for ‘fast’ growth of these species^34,54^. However, this did not influence stability because the same communities also had increased variation in productivity, likely because they were susceptible to drought. In the future, research should focus on how tree species richness, hydraulic diversity and community-weighted means of hydraulic traits affect population stability and potentially individual tree stability in addition to community stability. For instance, empirical work in forests found neutral^12^ or positive^7^ effects of species richness on population stability while studies in grasslands often reported a destabilizing effect of diversity on population stability^10^.

### Outlook

The frequency and severity of droughts and corresponding surges of tree mortality is dramatically increasing across the globe^31,32^. This situation is expected to worsen with intensifying climate change^1^, which threatens the climate mitigation potential of the world’s forests^3^. We show that the stability of forest community productivity along a 10-year observation period increases with tree species richness and that the key driver behind this diversity effect are the asynchronous growth dynamics of different tree species in hydraulically diverse communities. Importantly, stability did not compromise productivity. Instead, reduced temporal variation in productivity coincided with increased productivity in mixed-species tree communities. Hence, mixing tree species with a diversity of hydraulic strategies is likely a key management strategy to increase forest stability and their potential to mitigate the effects of climate change. Hydraulic traits may be used to select suitable tree species and design mixtures that stabilize productivity in an increasingly variable climate. Here, we examined the stability of young forest communities established as part of a large - scale biodiversity experiment. At the end of the observation period, tree height reached >10m in 25% of the experimental communities. It is conceivable that diversity effects on stability may strengthen as these stands mature, as indicated by the strengthening diversity effects on productivity^18^ and by results from an observational study that found stronger positive effects of species asynchrony on stability in old-growth than in secondary forests^25^. Our results extend research on forest stability from observational studies in relatively species-poor forests^7,12,13^ to species-rich subtropical tree communities growing under experimental conditions. This allowed establishing causality and avoiding confounding effects of environmental variation, major issues in observational studies. Stability increased consistently with tree species richness and did not plateau at low levels of tree species richness, which underlines the enormous potential of species richness to improve stability in many of our species-poor or mono-specific secondary and plantation forests around the world. This finding has important implications; contemporary forestry, and especially large-scale forest restoration initiatives^4^, like the Bonn Challenge, should focus on hydraulically diverse, mixed-species forests to enhance stability in a changing climate.

## Methods

### Study site and experimental design

In this study, we used data collected from the Tree Biodiversity–Ecosystem Functioning Experiment China (BEF-China, www.bef-china.com), located at Xingangshan, Dexing, Jiangxi (29° 08′–29° 11′N, 117° 90′–117°93′E). BEF-China^18,40^ is a large-scale tree biodiversity experiment that was established at two sites, A and B, each approximately 20 ha in size and planted in 2009 (site A) and 2010 (site B). The study sites are characterized by a subtropical, seasonal monsoon climate with hot and humid summers and dry and cool winters with a mean annual temperature of 16.7°C and mean annual precipitation of 1821mm^55^. The sites experienced strong inter-annual changes in climate-induced water availability during the 10-year observation period (Supplementary Fig. 9), with annual precipitation being more variable than temperature at our study sites^18^. The highly diverse native subtropical forests of the area are dominated by broadleaved mixed evergreen and deciduous tree species, sometimes interspersed with some conifers^40^. These forests are located in an area of overlap between tropical and temperate zones^56,57^, which makes them ideally suited to study diverse water use strategies and idiosyncratic species asynchrony as drivers of biodiversity–stability relationships. Furthermore, the region is densely populated and experiences frequent anthropogenic disturbances^56^, which makes the maintenance and improvement of the functioning of these forests important for the global ecosystem balance and restoration efforts.

The experiment covers a richness gradient ranging from 1–24 tree species. Communities have been assembled from a total pool of 40 native broadleaved evergreen and deciduous tree species (see Supplementary Table 3 for detailed species information). To ensure the representation of all species at each diversity level, mixture compositions were randomly allocated following a ‘broken-stick’ design^40^. In total 226,400 individual trees were planted on 566 plots^40^. In this study, we used data from six random extinction scenarios allocated to site A and B (three at each site) with a total of 396 plots and 158,400 planted trees^18^. Of these, we excluded 21 plots prior to our analysis due to failed establishment success, which left 375 plots (n=218 mixtures and n=157 monocultures) for our analysis. Each plot had a size of 25.8 × 25.8 m^2^ with 400 individual trees planted in 20 × 20 regular gridded positions (spacing 1.29m between trees). Tree positions and species compositions were randomly assigned to plots. More detailed information about the BEF-China experiment can be found in Huang *et al*. and Bruelheide *et al*.^18,40^.

### Tree data collection

Individual tree basal diameter at 5 cm above ground level (*gd*), tree height and species identity were measured annually from 2010 (site A) and 2011 (site B) onwards at the end of the growing season. To avoid edge effects, the central 12×12 trees were measured for each plot in the 4-, 8-, 16- and 24-species mixtures, while a smaller group of the central 6×6 trees was measured for monocultures and 2-species mixtures. Missing tree diameter and height values (in total 2% of census data) were imputed if the increment series was otherwise logical, i.e. *value*_*x*+1_ ≥ *value*_*x*−1_. To preserve climate-induced growth changes between years during imputation, we used a modelled site-specific rate of growth changes for each yearly step (*r*) based on complete increment series of trees with logical (i.e. with annual increases) and complete census data. A missing tree value was imputed as: (*v*_*x*+1_ − *v*_*x*−1_) * *r*_*x*_ + *v*_*x*−1_, where *v* is the *gd* or *height* measurement in a year and *r* the rate of change (see Supplementary Method 1 for details). Overall, we used annual data of 12,852 planted trees from 2010 to 2019 at site A and of 12,204 trees from 2011 to 2019 at site B to estimate community- and species-level productivity.

### Calculation of aboveground wood production

We used aboveground wood volume production as measure of community and species level productivity. First, annual aboveground wood volume per tree (awv, m^3^) was calculated with a fixed form factor of 0.5 (to account for the non-cylindrical shape of trees), which is an average value for the young subtropical trees in our experiment^21,58^; with

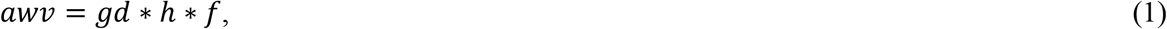

where *gd* is the basal area at measured tree ground diameter, *h* the measured tree height and *f* the form factor. Second, aboveground wood volume production (awp, m^3^ year^−1^) per tree and year was calculated as

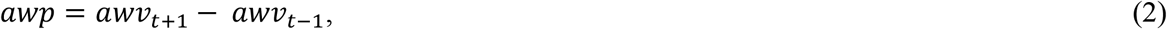

where *t* is an index for the year of measurement. Finally, *awv* and *awp* of all trees planted as part of the original design were summed per species and plot and scaled to 1 ha (based on the sampled subplot areas) to derive annual estimates of aboveground wood volume and volume production per species (AWV_s_, m^3^ ha^−1^; AWP_s_, m^3^ ha^−1^ year^−1^) and community (AWV, m^3^ ha^−1^; AWP, m^3^ ha^− 1^ year^−1^), referred to as species and community ‘productivity’. A value of 0 was used in case of species or plots with no alive tree individuals within individual years (note that completely failed plots were excluded from the analysis, see above). Our annual productivity estimates thus cover a complete series of forest growth over the course of 9 and 8 years for site A and B, respectively.

### Stability and asynchrony of production

The temporal stability^17^ of tree community productivity, hereafter ‘stability’, was calculated as the inverse of the coefficient of variation :

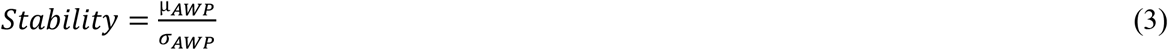

where µ_AWP_ is the temporal mean and σ_AWP_ the temporal standard deviation of annual plot productivity for our observation period (2010–2019 for site A and 2011–2019 for site B). Thus, any diversity effect that leads to overyielding (a higher productivity of mixtures vs monocultures) increases stability through increasing temporal mean productivity µ_AWP_. Conversely, any diversity effect that buffers variations in productivity against changing climatic conditions would increase stability through decreasing σ_AWP_^15^. We hypothesize here that asynchronous species growth dynamics to changing climatic conditions is the dominant mechanism that stabilizes young tree communities through lowering their productivity variance. To test this we calculated community-level species asynchrony using the species synchrony statistic φ^22^ as 1 – φ:

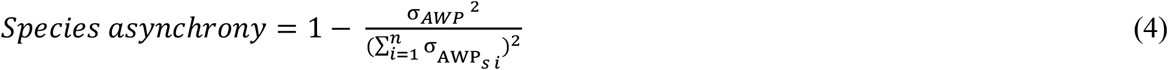

where σ_AWPs i_ is the temporal standard deviation of the annual productivity of species *i* in a plot of *n* species^7,59^. Thus, species asynchrony increases if the variance in individual species productivity increases relative to the variance in community productivity. Species asynchrony ranges from 0 (complete synchrony) to 1 (complete asynchrony) and is per definition 0 in monocultures as here variations in community productivity result from variations within a single species^7^. We expect here that species asynchrony increases stability through lowering the variation in community level productivity^7^. Young tree communities, as the ones examined here, show a strongly increasing productivity over time. As this age trend strongly masks annual variations in productivity, we removed it and calculated stability as temporal mean productivity divided by its detrended standard deviation. Similarly, species asynchrony was calculated based on detrended plot and species level productivity. Detrending was performed for each plot and species per plot through regressing annual productivity against time and then calculating the standard deviation based on the residuals of this regression following Craven *et al*. and Tilman *et al*.^8,10^ (see Supplementary Fig. 10 for a visualization of this approach).

### Trait gradients

Species employ different strategies to cope with climate induced water variability, which are likely related to a set of (hydraulic) functional traits (Anderegg *et al*.^39^ and citations within). We assembled species-specific hydraulic trait data related to stomatal control and drought tolerance that was measured within the experiment (Supplementary Table 1; refs.^37,38^). Trait data were subjected to a principal component analysis (PCA). The first and second axis partitioned the hydraulic traits into two orthogonal trait gradients related to stomatal control (PC1) and drought tolerance (PC2) (Supplementary Fig. 1). Based on physiological and morphological leaf traits, we classified species as water spenders if they decrease their stomatal conductance only at high levels of water pressure deficit, and as water savers, if they already decrease stomatal conductance at low water pressure deficits and have leaves characterized by high stomatal density. We used the water potential at which 50% of xylem conductivity is lost (Ψ_50_) as key physiological trait to quantify a species drought tolerance^2^. Higher values of Ψ_50_ (i.e. lower absolute values Ψ_50_) indicate a higher susceptibility to drought-induced xylem cavitation. We also included specific leaf area, leaf toughness and carbon to nitrogen ratio as classic traits of the leaf economics spectrum (LES^34^) in our analysis, which are associated with a species drought tolerance^37^, to foster the still limited understanding of trait syndromes that govern forest responses to climatic stress^36^. We used trait data from 39 out of the 40 planted species (*Castanopsis carlesii* was excluded due to complete establishment failure) and imputed two missing trait values (Ψ_50_ and stomatal density) for one out of these 39 species (*Quercus phillyreoides*) with predicted mean value matching with 500 runs using the R package mice^60^. PCA was performed with the *rda* function in the vegan package version 2.5-6^61^.

### Quantifying hydraulic diversity and community means

We used the scores of the first and second PCA axis (Supplementary Fig. 1) as measure of the species stomatal control and drought tolerance strategies within each community. Functional diversity in traits associated with water spending vs water saving stomatal behaviour (hereafter ‘functional diversity of stomatal control’) and functional diversity of drought tolerance was calculated with the ‘FD’ package as abundance-weighted functional dispersion^62,63^ using temporal mean species wood volume per plot as measure of species abundance. Functional dispersion measures the mean distance of species along each trait gradient^62^ and thus represents the complementarity in hydraulic strategies of co-occurring species within each community. We calculated community-weighted mean (CWM) trait values for both gradients, hereafter called ‘CWM of stomatal control ‘ and ‘CWM of drought tolerance’ using temporal mean species wood volume per plot as measure of species abundance.

### Modelling framework and statistical analysis

First, we analysed direct relationships between stability, its hypothesized drivers and relationships between these drivers. Specifically, we used linear mixed-effect models (LMM) to test for bivariate relationships between species richness, species asynchrony, functional diversity of stomatal control, functional diversity of drought tolerance, CWM of stomatal control and the CWM of drought tolerance. We also tested the effect of species richness and hydraulic diversity on species asynchrony. LMM were fit with the nlme package version 3.1-144^64^ to allow for the specification of variance functions with a significance level of α=0.05. Confidence intervals (95%) of LMM effects were computed with the ggeffects package^65^. Tree species richness was log_2_ transformed in all models. As the two sites were planted one year apart, we tested for a potential age effect and other site-specific influences on the biodiversity–stability relationship through including site and its interaction with species richness as fixed effect. Diversity effects on stability did not differ between sites (P=0.46 for the interaction). We therefore accounted for site and other aspects of our experimental design through a nested random effect structure of site, species composition and arrangement of plots within quadrants (see Huang *et al*.^18^). Model assumptions were visually checked for independence and homogeneity of variance through examining model residuals and for normal distribution with quantile-quantile plots. For all response variables we tested the inclusion of an exponential variance structure^64^ to model heteroscedasticity (parsimony evaluated via AIC) and a log/square-root transformation to normalize residuals. As results did not differ for any bivariate relationship, we present only the models without variance function or transformation of response variables.

Second, we developed a hypothesis driven structural equation model (SEM) framework to disentangle direct and indirect drivers of stability based on *a priori* knowledge of mechanisms driving biodiversity–stability relationships (Supplementary Fig. 7). We explored whether the data supported our first and second hypothesis through including indirect pathways that tested for effects of the multiple diversity facets species richness, functional diversity of stomatal control and functional diversity of drought tolerance on stability through effects mediated via species asynchrony. We also included direct pathways from these diversity facets to stability, to test for mechanisms not mediated by species asynchrony such as performance enhancing effects that increase temporal mean productivity in mixtures^8,14,18^. To test for the effects of community trait means we included direct pathways from the CWM of stomatal control and the CWM of drought tolerance to stability^8,39^. As the experimental manipulation of species richness may directly affect the functional diversity of a community^40^, we included pathways from species richness to functional diversity of stomatal control and functional diversity of drought tolerance. Piecewise SEMs^41^ were used to test the support for and relative importance of these hypothesized pathways. To understand whether diversity effects on stability result from overyielding (increased µ_AWP_), a buffered variation (decreased σ_AWP_) or both, we fit a separate SEM with these two components of our temporal stability measure as response. In this second SEM, we tested all hypothesized effects of diversity on stability for each of its two components (Supplementary Fig. 8).

Global model fit was assessed via Fisher’s C statistic (P>0.05). We assessed the independence of variables and included partial, non-directional correlations if these improved model fit based on tests of directed separations (P<0.05 for violation of independence claims)^41^. For each SEM we calculated standardized path coefficients, which allowed us to compare the strength of paths within and among models and of indirect pathways (calculated as product of the coefficients along the path)^41^. We fitted individual pathways with LMM using the same random structure and model evaluation as for our analysis of bivariate relationships detailed above. In all SEMs stability, species asynchrony, the temporal mean (µ_AWP_) and the temporal standard deviation of productivity (σ_AWP_) were square-root transformed to best meet model assumptions. Our analysis focuses on the role of species asynchrony and hydraulic diversity as drivers of biodiversity–stability relationships. As species asynchrony and functional diversity in monocultures are per definition 0, we analysed their effects within 2-, 4-, 8-, 16- and 24-species mixtures only to avoid many observations without variation. Alternative models including monocultures yielded the same results for effects reported here (Supplementary Figs. 6, 11–12). To further test the sensitivity of our models, we ran alternative SEMs without response transformation but with an exponential variance structure for log2 species richness. These yielded the same results (Supplementary Figs. 13–14). SEMs had low variance inflation (Variance Inflation Factor < 5, a conservative threshold choice^66^). All analyses were performed in R 3.6.2^67^.

## Supporting information

Supplementary material_Schnabel_et_al

## Acknowledgements

We thank local workers for their invaluable help in the field. In particular, we acknowledge the work of Wenzel Kröber on hydraulic trait measurements. This study was supported by the German Research Foundation (DFG FOR 891), the National Key Research and Development Program of China (2017YFA0605103) and the Strategic Priority Research Program of the Chinese Academy of Sciences (XDB31000000). K.E.B. was supported by the German Centre for Integrative Biodiversity Research (iDiv)’s flexible pool initiative (grant #34600900). H.B. acknowledges the DFG support for trait measurements (DFG BR 1698/9-2). B.S. was supported by the URPP Global Change and Biodiversity of the University of Zurich. F.S. was supported by the International Research Training Group TreeDì funded by the Deutsche Forschungsgemeinschaft (DFG, German Research Foundation) – 319936945/GRK2324 and the University of Chinese Academy of Sciences (UCAS).

## Author contributions

F.S. and X.L. are co-first authors. H.B., W.H., B.S., Z.T., B.Y., J.B., G.v.O., K.M. and C.W. designed the project; F.S., X.L., K. E.B., J.A.S., J.B. and C.W. conceived the idea for the manuscript; X.L., M.K., G.v.O., H.B., F.J.B., A.F., S.L., C.T.P. and F.S. collected and compiled data; F.S. analysed and interpreted the data and wrote the manuscript with support from X.L., K.E.B. and C.W.; F.S. and K.E.B. created figures; All authors discussed the results and contributed substantially to revisions.

## Data availability statement

Data supporting the findings of this study have been deposited on the BEF-China project database (https://data.botanik.uni-halle.de/bef-china/datasets/634) and are available upon reasonable request from the corresponding authors.

