## Supplementary material_Schnabel_et_al for "Hydraulic diversity stabilizes productivity in a large-scale subtropical tree biodiversity experiment"

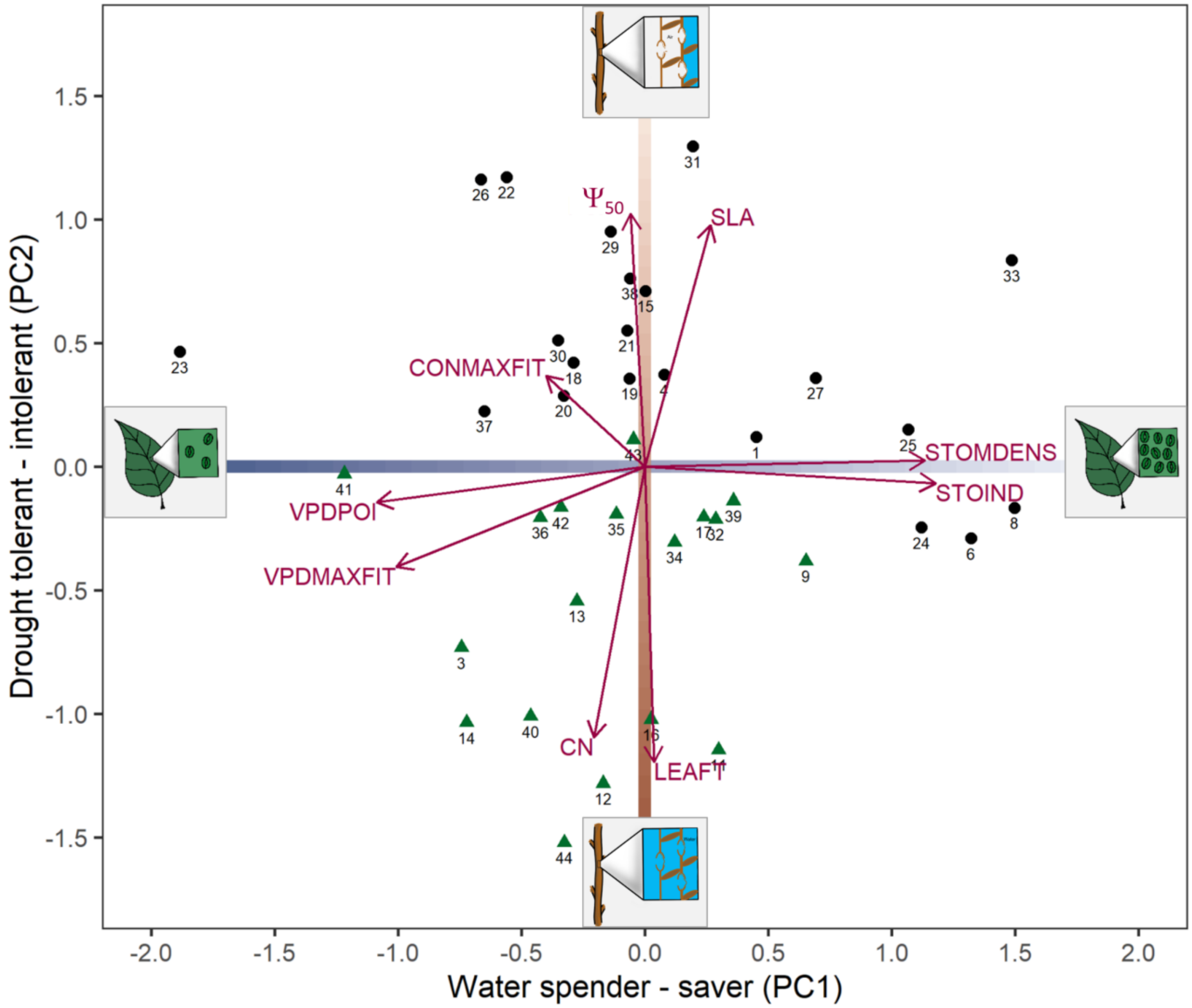


**Supplementary Figure 1** Principal component analysis (PCA) biplot for the trait based definition of two hydraulic trait gradients: (1) A gradient of stomata control that characterizes species as 'water spenders' if they down-regulate their stomatal conductance only at high levels of measured water pressure deficit (high VPDPOI and VPDMAXFIT), while 'water savers' are species that already down-regulate stomatal conductance at low water pressure deficits and have leaves characterized by a high a stomata density (high STODENS, STOIND; see Supplementary Table 1 for a trait description). (2) A gradient of drought tolerance based on the water potential at which 50% of xylem conductivity is lost ($\Psi_{50}$) as key physiological trait that expresses a species resistance to water stress[1], with less negative values of $\Psi_{50}$ indicating a higher susceptibility to drought induced cavitation. The sketches schematically illustrate the trait gradients: water-spending vs water-saving stomatal control (few versus abundant stomata) and drought tolerance (high versus low cavitation resistance). Note, that classic traits of the leaf economic spectrum (LES[2]) are associated with drought tolerance in that drought-tolerant species also have traits commonly used to ascribe a conservative resource use strategy (tough leaves (LEAFT) and a high C/N ration (CN)) while drought intolerance is associated to a high resource acquisitiveness (high SLA)[3–5]. All traits (Supplementary Table 1) were measured on site and used to calculate species level mean trait values, see refs.[3,6] for details. Species identity is shown as species code; see Supplementary Table 3 for a detailed species list. Deciduous and evergreen species are coloured as black dots and green triangles, respectively. PCA was performed with the *rda* function in the vegan package version 2.5-6[7]. Varimax rotated principal components (base R version 3.6.2[8]) were used in the analysis to achieve a good alignment of the two orthogonal trait gradients with the first and second PCA axis.

**Supplementary Table 1** Drought tolerance and stomatal control traits as well as traits of the leaf economic spectrum (LES[2]) used in this study.

| Acronym | Trait description | Unit |
|---|---|---|
| $\Psi_{50}$ | Water potential at which 50% initial conductivity is lost | MPa |
| CONMAXFIT | Modelled maximum stomata conductance | Non-dimensional |
| VPDMAXFIT | Water pressure deficit (VPD) at CONMAXFIT | hPA |
| VPDPOI | VPD at the point of inflection of modelled stomatal conductance | hPA |
| SLA | Specific leaf area | $m^2$ $kg^{-1}$ |
| LEAFT | Leaf toughness | N $mm^{-1}$ |
| CN | Carbon to nitrogen ratio | Ratio |
| STOMDENS | Stomata density | 1 $mm^{-2}$ |
| STOIND | Product of STODENS and stomata size in $\mu m^2$ | ratio |

Note: All traits were measured on site and used to calculate species level mean trait values by refs.[3,6]. See these studies for a detailed explication of the selected traits and Supplementary Figure 1 for additional information. In brief, CONMAXFIT is the modelled maximum stomata conductance, which allows calculating the following key physiological traits of stomata control relevant under water shortage: VPDMAXFIT is the water pressure deficit (VPD) at CONMAXFIT and VPDPOI the VPD at the second point of inflection of a modelled curve of stomatal conductance regressed against VPD, i.e. a measure of stomata sensitivity[6]. Morphological stomata traits are stomata density and stomata index, the product of stomata size and stomata density.

**Supplementary Table 2** Mixed-effect models exploring bivariate relationships between stability, species asynchrony and different facets of hydraulic diversity and community hydraulic means.

| Response | Fixed Effects | ddf | t | P-value | n | $R^2_m$ | $R^2_c$ |
|---|---|---|---|---|---|---|---|
| **All plots** | | | | | | | |
| Stability | **Species richness** | 132 | 3.98 | **0.000** | 375 | 0.06 | 0.26 |
| **Mixtures only** | | | | | | | |
| Stability | **Species richness** | 87 | 3.72 | **0.000** | 218 | 0.08 | 0.22 |
| Stability | **Species asynchrony** | 95 | 10.13 | **0.000** | 218 | 0.34 | 0.54 |
| Stability | **FD stomatal control** | 95 | 1.92 | **0.058** | 218 | 0.02 | 0.22 |
| Stability | FD drought tolerance | 95 | 1.12 | 0.270 | 218 | 0.01 | 0.21 |
| Stability | CWM stomatal control | 95 | -0.45 | 0.652 | 218 | 0.00 | 0.21 |
| Stability | CWM drought tolerance | 95 | -0.15 | 0.880 | 218 | 0.00 | 0.21 |
| Species asynchrony | **Species richness** | 87 | 9.53 | **0.000** | 218 | 0.38 | 0.49 |
| Species asynchrony | **FD stomatal control** | 95 | 5.29 | **0.000** | 218 | 0.16 | 0.49 |
| Species asynchrony | **FD drought tolerance** | 95 | 5.84 | **0.000** | 218 | 0.17 | 0.43 |

Note: Significant fixed effects ($P<0.05$) printed in bold. All models have the same nested random effect structure of species composition and plot arrangement nested within site (see ref.[9]) and were fit with residual maximum likelihood (REML) estimation using the package nlme[10]. Hydraulic diversity was quantified as functional diversity of stomatal control (FD stomatal control) and functional diversity of drought tolerance (FD drought tolerance) and calculated as abundance-weighted functional dispersion. Community hydraulic means were quantified as community-weighted mean of stomatal control (CWM stomatal control) and drought tolerance (CWM drought tolerance). Data is based on a long, experimental species richness gradient with mixtures of 2, 4, 8, 16 and 24 tree species. ddf are the denominator degrees of freedom; t the ratio between the estimate and its standard error; p-value from a t-distribution; n the number of plots; Marginal $R^2$ values ($R^2_m$) show the variance explained by fixed effects and conditional ($R^2_c$) values the variance explained by fixed and random effects[11].

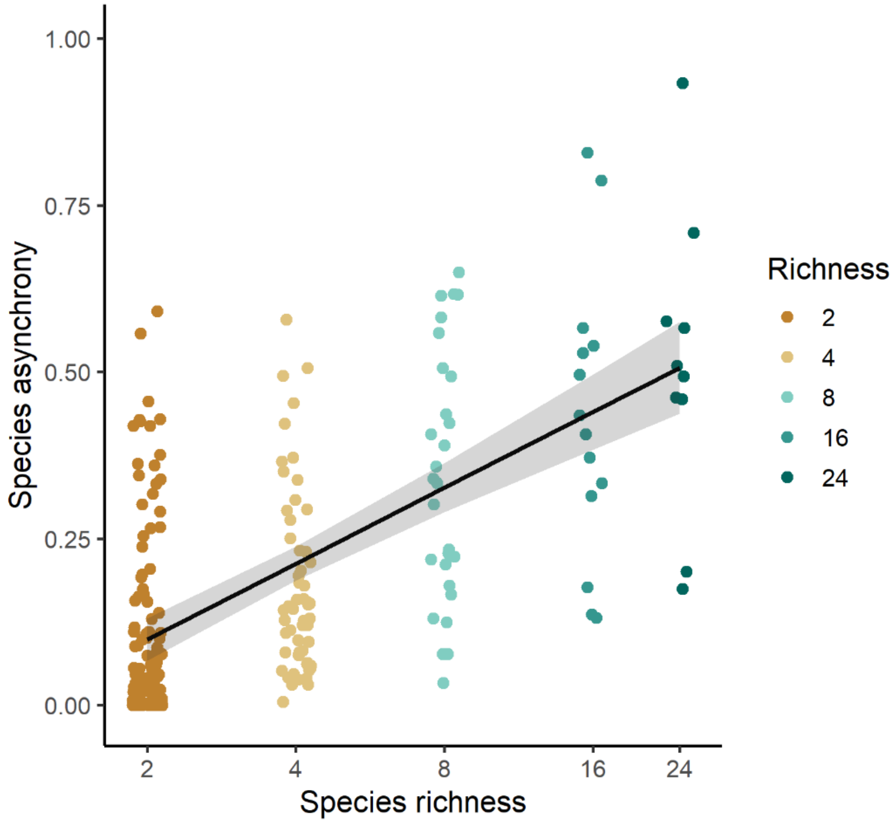


**Supplementary Figure 2** Direct effects of species richness on species asynchrony. The black line is the fit of a linear mixed-effect model that shows significant increases ($P<0.001$) in species asynchrony with species richness in mixtures. Species asynchrony ranges from 0 to 1. 0 represents complete synchrony and 1 represents complete asynchrony. Grey bands represent a 95% confidence interval. See Supplementary Table 2 for details on the fitted model.

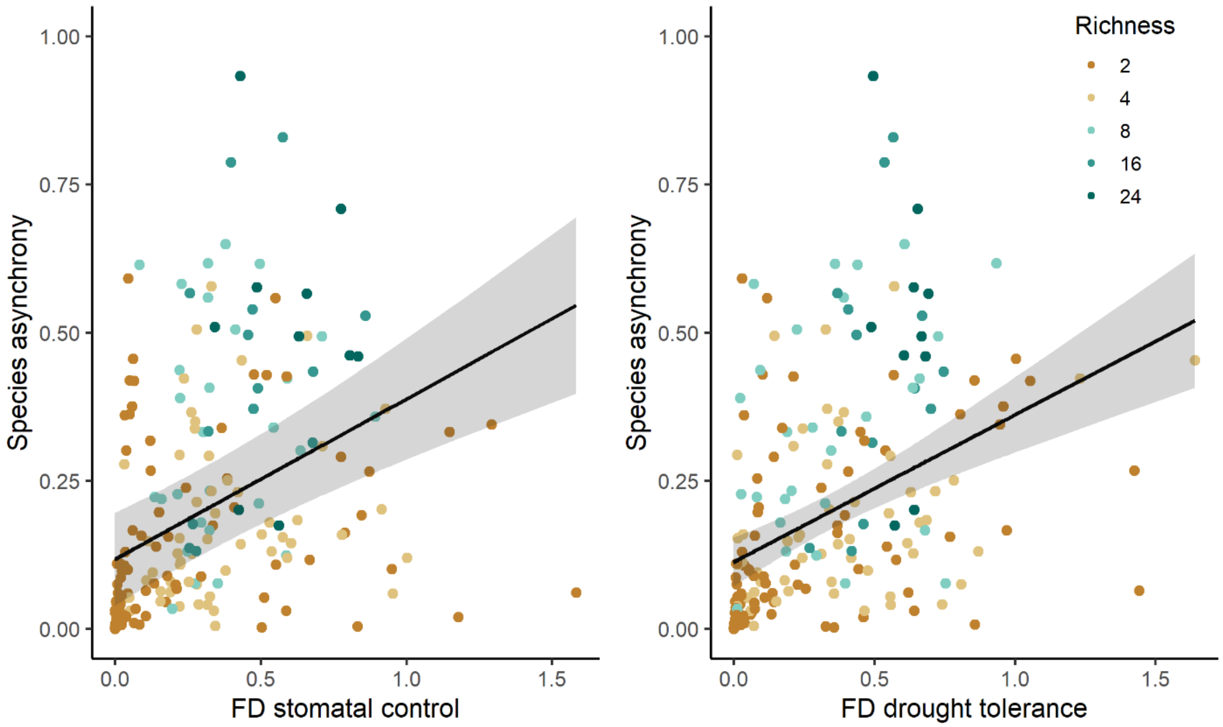


**Supplementary Figure 3** Direct effects of hydraulic diversity on species asynchrony. Lines are linear mixed-effect model fits that show (a) significant increases in species asynchrony with functional diversity of stomatal control (FD stomatal control; $P<0.001$) and (b) significant increases in species asynchrony with functional diversity of drought tolerance (FD drought tolerance; $P<0.001$) in mixtures. Species asynchrony ranges from 0 to 1. 0 represents complete synchrony and 1 represents complete asynchrony. Functional diversity calculated as abundance-weighted functional dispersion. Grey bands represent a 95% confidence interval. See Supplementary Table 2 for details on the fitted models.

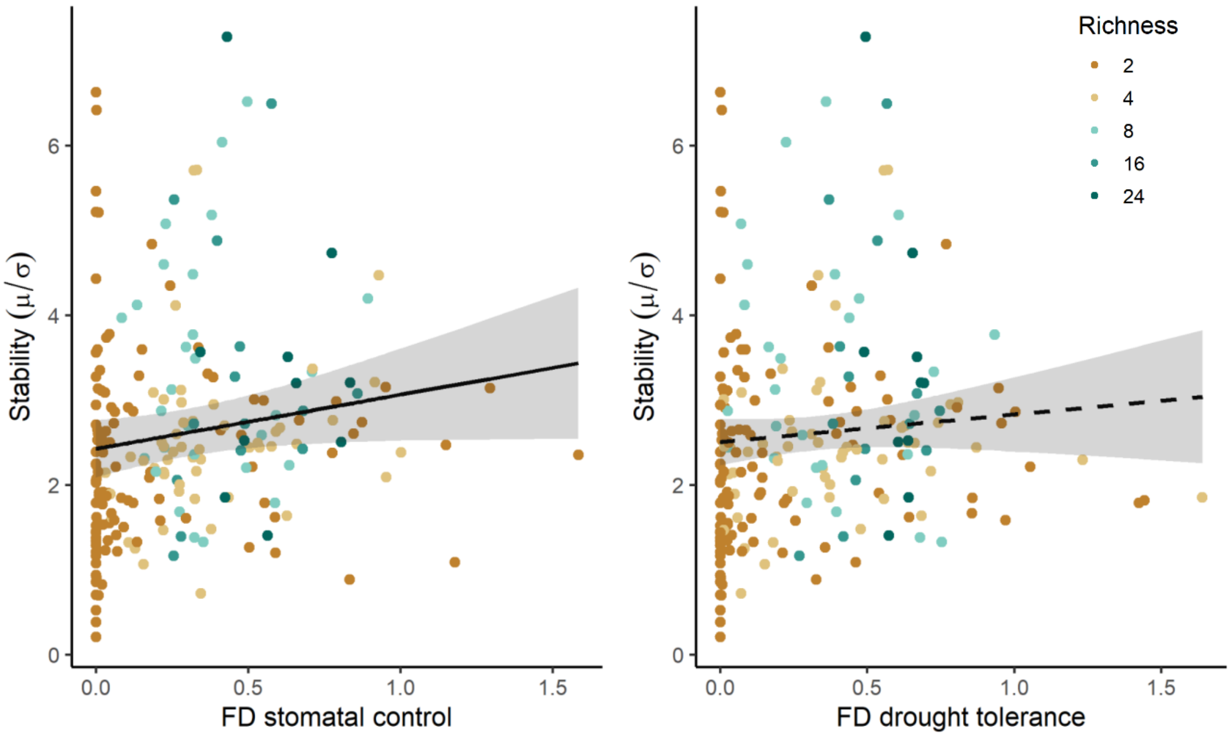


**Supplementary Figure 4** Direct effects of hydraulic diversity on stability. Lines are linear mixed-effect model fits that show (a) a marginally significant increase in stability with functional diversity of stomatal control (FD stomatal control; P=0.058) and (b) a non-significant relationship of stability with functional diversity of drought tolerance (FD drought tolerance; P=0.27) in mixtures. Species asynchrony ranges from 0 to 1. 0 represents complete synchrony and 1 represents complete asynchrony. Functional diversity calculated as abundance-weighted functional dispersion. Grey bands represent a 95% confidence interval. See Supplementary Table 2 for details on the fitted models.

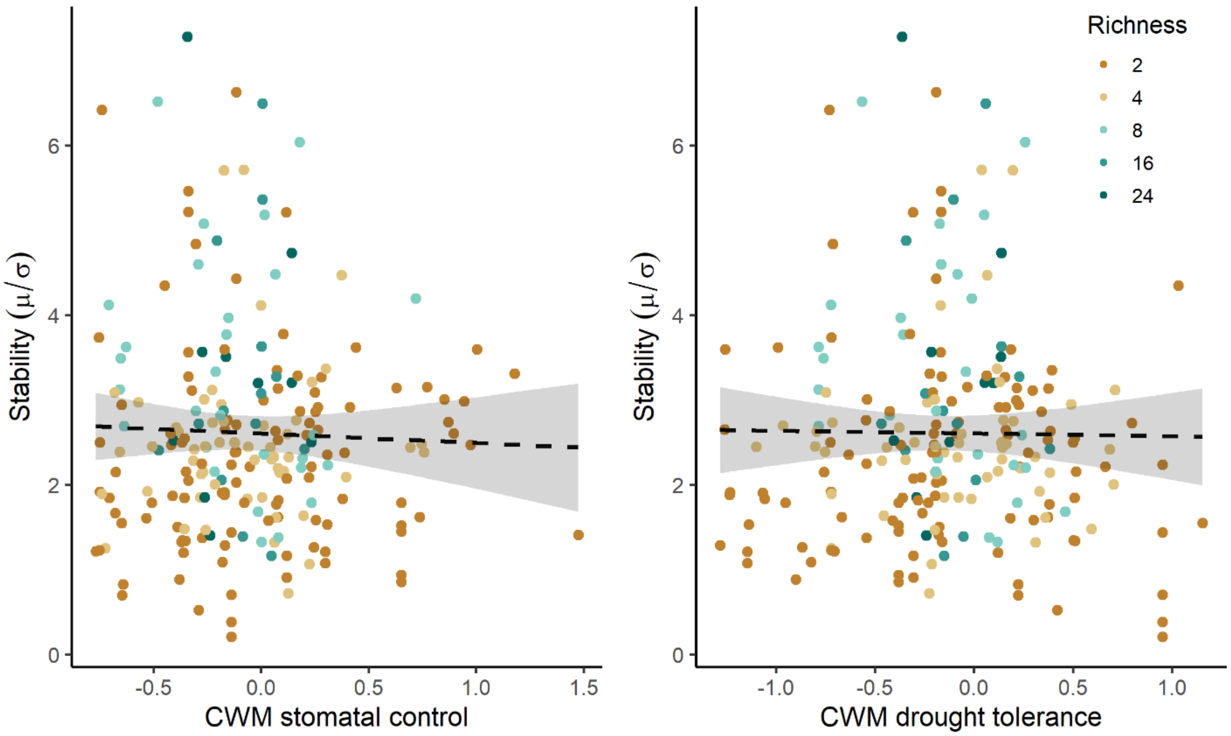


**Supplementary Figure 5** Direct effects of community-weighted hydraulic means (CWMs) on stability. Lines are linear mixed-effect model fits that show (a) a non-significant relationship of stability with the CWM of stomatal control (CWM stomatal control; P=0.65) and (b) a non-significant relationship of stability with the CWM of drought tolerance (CWM drought tolerance; P=0.88) in mixtures. Grey bands represent a 95% confidence interval. See Supplementary Table 2 for details on the fitted models.

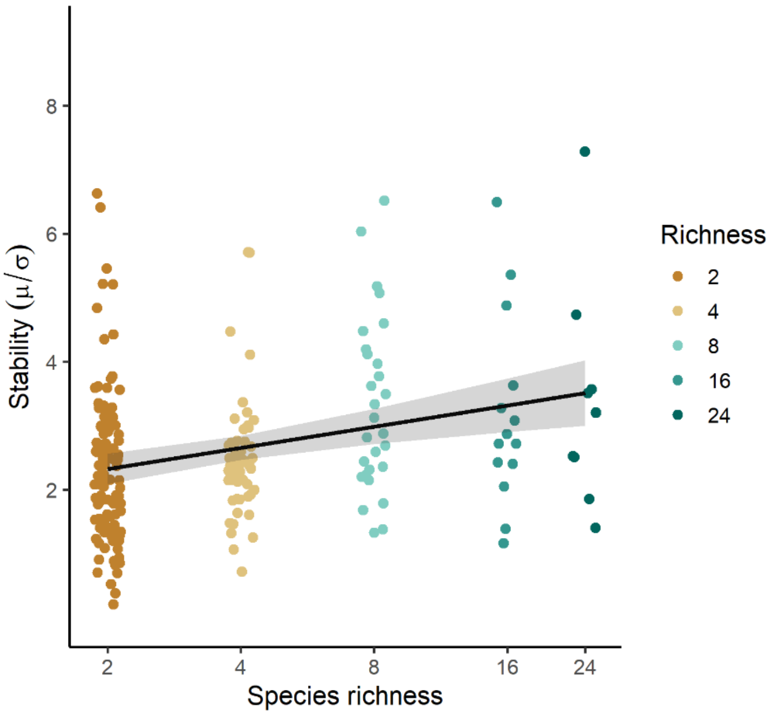


**Supplementary Figure 6** Effects of species richness on stability within mixtures. The black line is the fit of a linear mixed-effect model (LMM) that shows a significant increase ($P<0.001$) in stability with species richness along a planted diversity gradient ranging from 2-species mixtures up to 24-species mixtures (n=218 plots). The shown model has a similar model fit compared to the model shown in Fig. 2 which was fit to data from all plots, including monocultures (n=375 plots; see Supplementary Table 2 for a detailed model comparison).

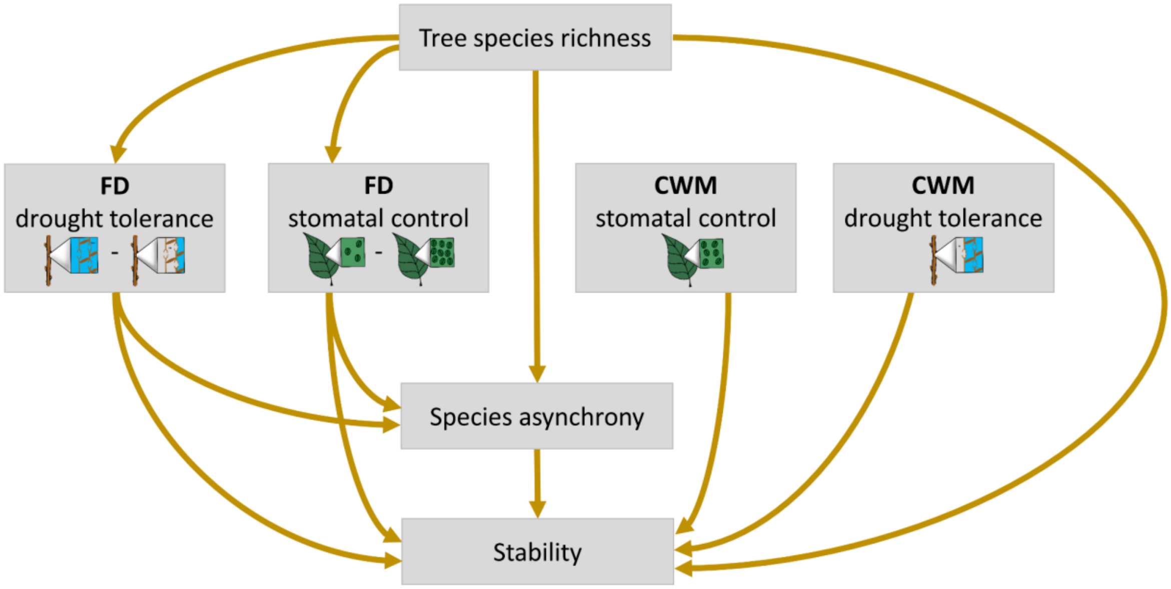


**Supplementary Figure 7** Hypotheses driven framework of the direct and indirect drivers of stability in mixed-species tree communities. Directional arrows represent hypothesized pathways. The causal model predicts positive effects of diversity - species richness, functional diversity of stomatal control and functional diversity of drought tolerance - on stability through effects mediated via species asynchrony. We also included direct pathways from these diversity facets to stability, to test for mechanisms not mediated by species asynchrony such as performance enhancing effects that increase temporal mean productivity in mixtures[9,12–14]. To test for community trait mean effects we included direct pathways from the CWM of stomatal control and the CWM of drought tolerance to stability[13,15]. Finally, we included pathways from species richness to functional diversity of stomatal control and functional diversity of drought tolerance as the experimental manipulation of species richness in our experiment may directly affect functional diversity[16]. The sketches schematically illustrate the trait gradients: water-spending vs water-saving stomatal control (few versus abundant stomata) and drought tolerance (high versus low cavitation resistance). This framework was tested with data from experimental tree communities from the BEF-China experiment[9,16], that span a long gradient of planted tree species richness with mixtures of up to 24 different tree species using piecewise structural equation models[17] (SEMs).

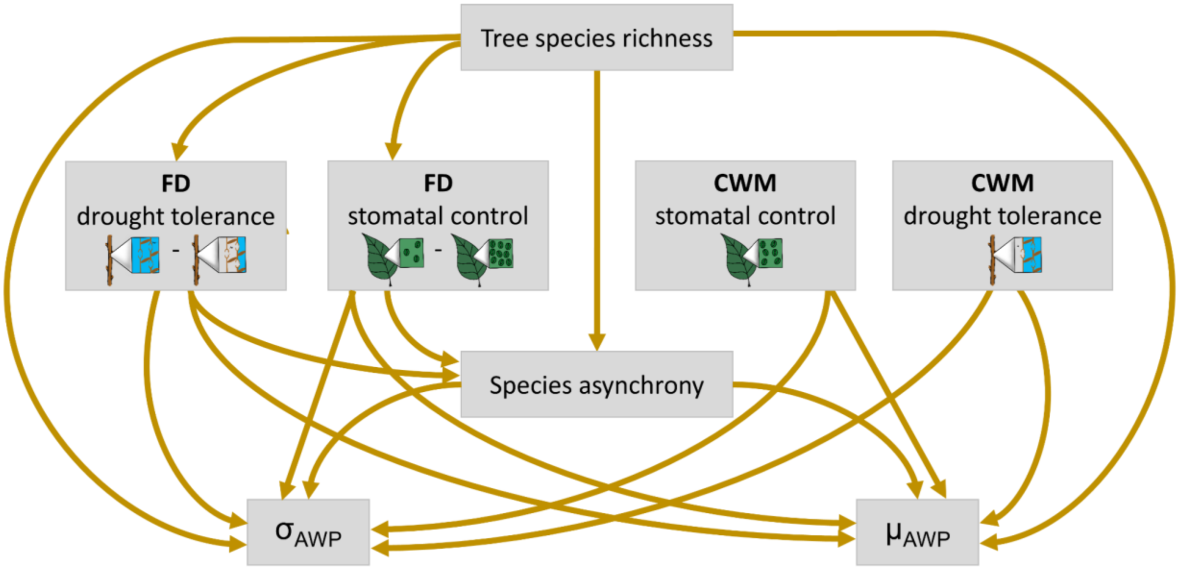


**Supplementary Figure 8** Hypotheses driven framework for the partitioning of the direct and indirect effects of species richness, hydraulic diversity and community hydraulic means into overyielding and variance buffering effects of diversity. Directional arrows represent hypothesized pathways. The framework separates the hypothesized effects of diversity on stability (Supplementary Figure 7) into effects on the temporal mean ($\mu_{AWP}$) and the temporal standard deviation of productivity ($\sigma_{AWP}$). Increases in $\mu_{AWP}$ would enhance stability through overyielding – a higher productivity in mixtures vs monocultures - and decreases in $\sigma_{AWP}$ would enhance stability through buffered variations in productivity. All drivers hypothesized to influence stability (Supplementary Figure 7) – species richness, functional diversity of stomatal control (FD stomatal control), functional diversity of drought tolerance (FD drought tolerance), the CWM of stomatal control (CWM stomatal control), the CWM of drought tolerance (CWM drought tolerance) and species asynchrony – are partitioned here into their effects on $\mu_{AWP}$ and $\sigma_{AWP}$. The sketches schematically illustrate the trait gradients: water-spending vs water-saving stomatal control (few versus abundant stomata) and drought tolerance (high versus low cavitation resistance). This framework was tested with data from experimental tree communities from the BEF-China experiment[9,16], that span a long gradient of planted tree species richness with mixtures of up to 24 different tree species using piecewise structural equation models[17] (SEMs).

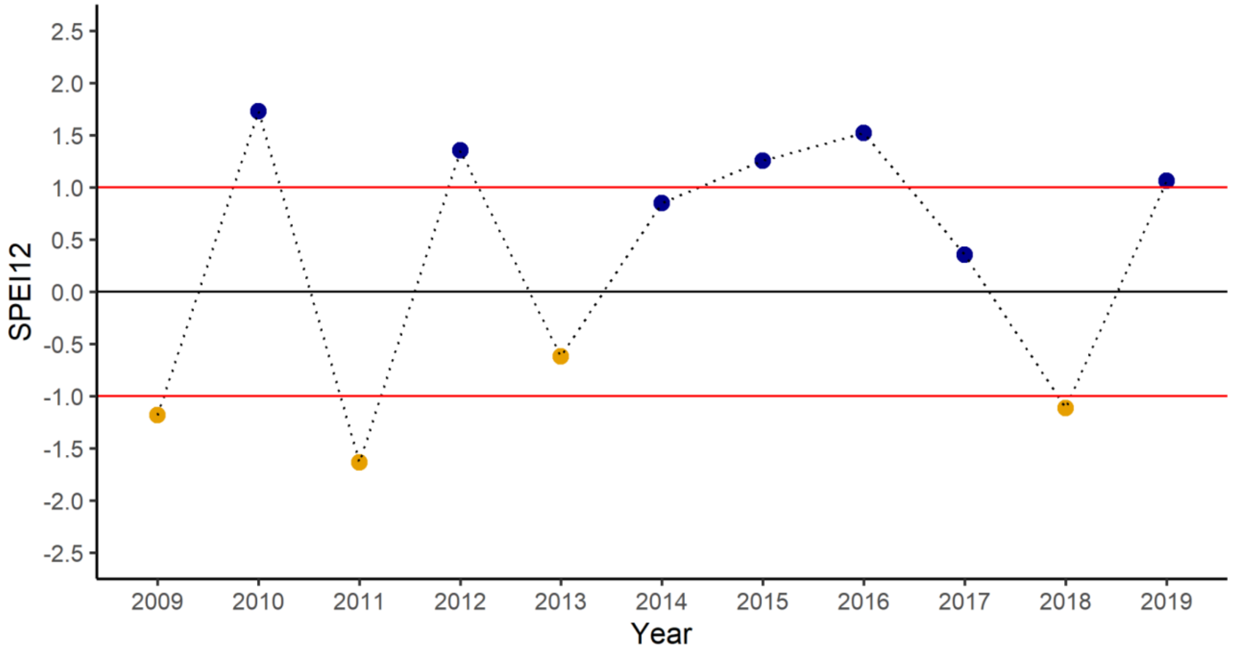


**Supplementary Figure 9** Annual values of the Standardised Precipitation-Evapotranspiration Index (SPEI). The drought index[18] captures the annual standardized water balance (precipitation minus potential evapotranspiration) and its variation during our study period. Negative values indicate climatic water deficits (brown coloured points) and positive values a water surplus (blue coloured points). Years with values above 1 or below -1 can be considered exceptionally wet and dry, respectively. The SPEI index was calculated based on monthly resolved precipitation and potential evapotranspiration data (CRU TS 4.04[19]) and is expressed here as annual water balance between two tree census intervals (SPEI12, water balance from September to October of the preceding year). The SPEI index was calculated with the SPEI package[20] in R using climate data from 1901-2019 as a reference period.

**Supplementary Table 3** List of the 40 planted broadleaved evergreen and deciduous tree species.

| Species names | Family | Species code | Leaf habit | Site |
|---|---|---|---|---|
| *Acer davidii* | *Sapindaceae* | 27 | D | A |
| *Ailanthus altissima* | *Simaroubaceae* | 29 | D | B |
| *Alniphyllum fortunei* | *Styracaceae* | 30 | D | B |
| *Betula luminifera* | *Betulaceae* | 31 | D | B |
| *Castanea henryi* | *Fagaceae* | 1 | D | A |
| *Castanopsis carlesii* | *Fagaceae* | 10 | E | A |
| *Castanopsis eyrei* | *Fagaceae* | 13 | E | AB |
| *Castanopsis fargesii* | *Fagaceae* | 32 | E | B |
| *Castanopsis sclerophylla* | *Fagaceae* | 14 | E | AB |
| *Celtis biondii* | *Cannabaceae* | 33 | D | B |
| *Choerospondias axillaris* | *Anacardiaceae* | 4 | D | A |
| *Cinnamomum camphora* | *Lauraceae* | 17 | E | AB |
| *Cyclobalanopsis glauca* | *Fagaceae* | 11 | E | AB |
| *Cyclobalanopsis myrsinaefolia* | *Fagaceae* | 9 | E | A |
| *Daphniphyllum oldhamii* | *Daphniphyllaceae* | 16 | E | AB |
| *Diospyros japonica* | *Ebenaceae* | 15 | D | AB |
| *Elaeocarpus chinensis* | *Elaeocarpaceae* | 34 | E | B |
| *Elaeocarpus glabripetalus* | *Elaeocarpaceae* | 35 | E | B |
| *Elaeocarpus japonicus* | *Elaeocarpaceae* | 36 | E | B |
| *Idesia polycarpa* | *Salicaceae* | 37 | D | B |
| *Koelreuteria bipinnata* | *Sapindaceae* | 18 | D | A |
| *Liquidambar formosana* | *Altingiaceae* | 6 | D | A |
| *Lithocarpus glaber* | *Fagaceae* | 12 | E | AB |
| *Machilus grijsii* | *Lauraceae* | 39 | E | B |
| *Machilus leptophylla* | *Lauraceae* | 41 | E | B |
| *Machilus thunbergii* | *Lauraceae* | 40 | E | B |
| *Manglietia fordiana* | *Magnoliaceae* | 42 | E | B |
| *Melia azedarach* | *Meliaceae* | 26 | D | A |
| *Meliosma flexuosa* | *Sabiaceae* | 38 | D | B |
| *Nyssa sinensis* | *Cornaceae* | 20 | D | A |
| *Phoebe bournei* | *Lauraceae* | 43 | E | B |
| *Quercus acutissima* | *Fagaceae* | 25 | D | A |
| *Quercus fabri* | *Fagaceae* | 24 | D | A |
| *Quercus phillyreoides* | *Fagaceae* | 44 | E | B |
| *Quercus serrata* | *Fagaceae* | 8 | D | A |
| *Rhus chinensis* | *Anacardiaceae* | 23 | D | A |
| *Sapindus saponaria* | *Sapindaceae* | 19 | D | A |
| *Triadica cochinchinensis* | *Euphorbiaceae* | 22 | D | A |
| *Triadica sebifera* | *Euphorbiaceae* | 21 | D | A |
| *Schima superba* | *Theaceae* | 3 | E | AB |

Note: Shown are species and family names, species identity codes, leaf habit (E, evergreen; D, Deciduous) and the site at which the species were planted. For more details on the experimental design see refs.[9,16].

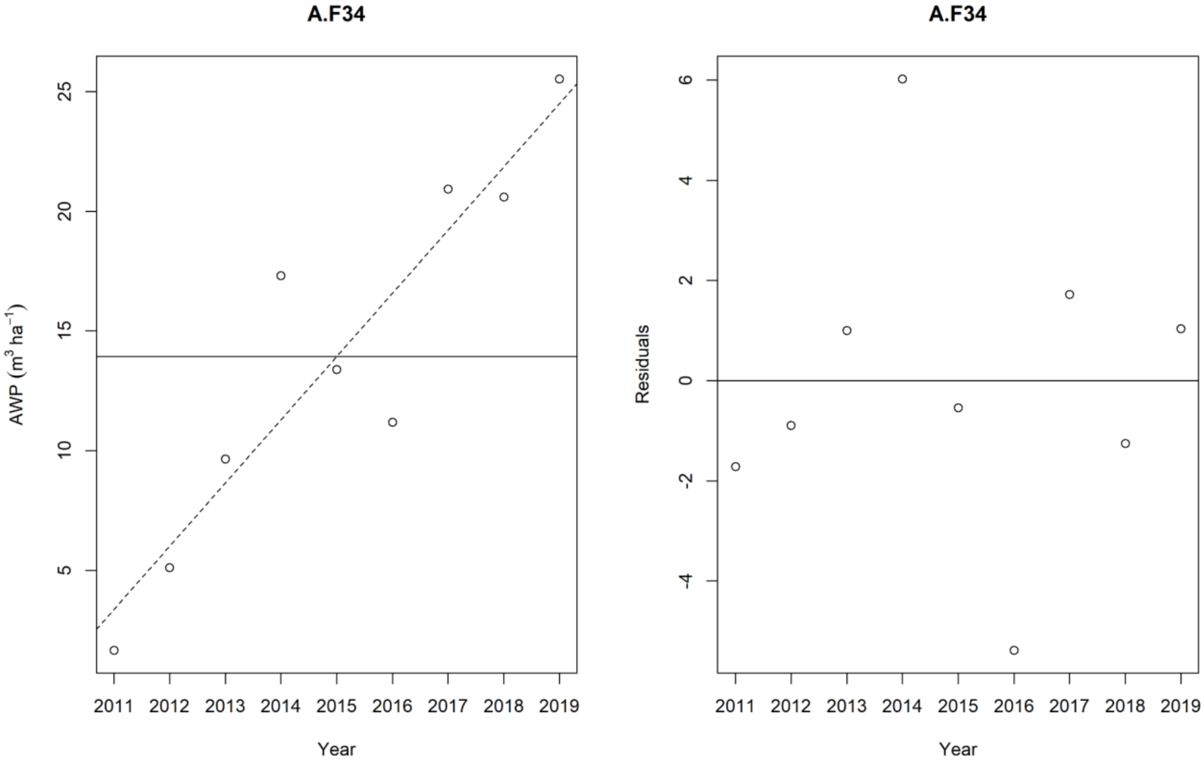


**Supplementary Figure 10** Example for the calculation of detrended stability. (a) Shows annual aboveground wood volume productivity (AWP, from plot F34 on site A) regressed against time. (b) The residuals of this regression represent the annual variation in productivity without a directional stand development trend. The standard deviation of these residuals gives the detrended standard deviation that was subsequently used to calculate detrended stability as temporal mean productivity ($\mu_{AWP}$) divided by its detrended standard deviation ($\sigma_{AWP}$).

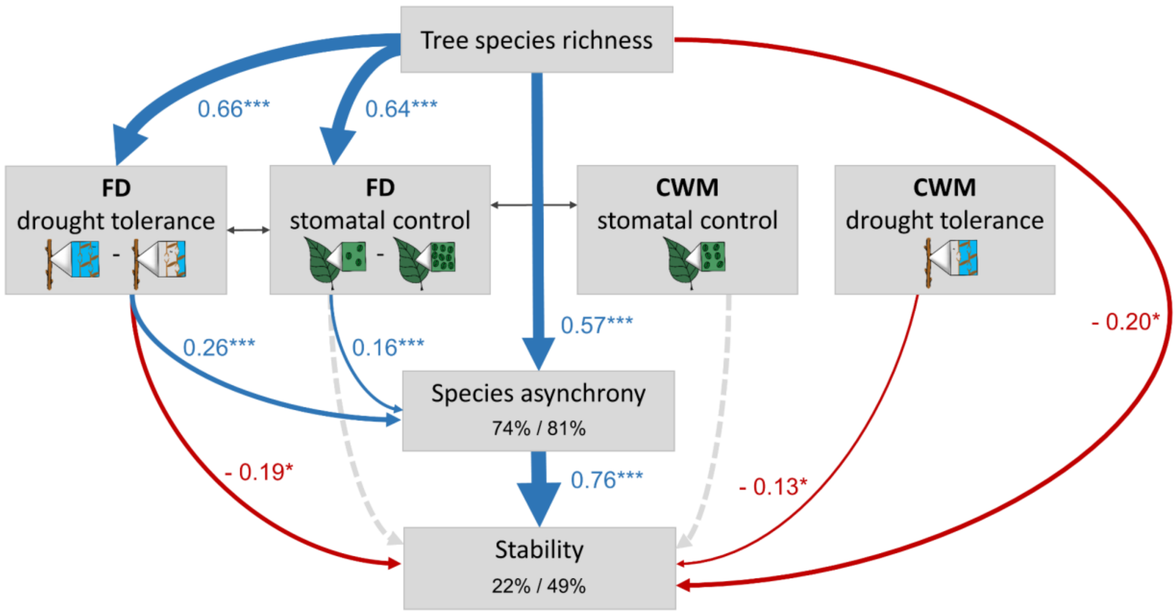


**Supplementary Figure 11** Direct and indirect effects of species richness, hydraulic diversity and community hydraulic means on stability. The structural equation model (SEM) is fit to data including monocultures and tests the effects of tree species richness, functional diversity of stomatal control (FD stomatal control) and functional diversity of drought tolerance (FD drought tolerance) on species asynchrony and stability. The sketches schematically illustrate the trait gradients: water-spending vs water-saving stomatal control (few versus abundant stomata) and drought tolerance (high versus low cavitation resistance). Functional diversity was calculated as abundance-weighted functional dispersion. Effects of community-weighted mean (CWM) traits are explored through testing the effect of the CWM of stomatal control (CWM stomatal control) and the CWM of drought tolerance (CWM drought tolerance) on stability. The SEM fit the data well (Fisher's C=10.7, P=0.22, d.f.=8, n=375 plots). Data is based on a long, experimental species richness gradient ranging from monocultures to mixtures of 24 tree species. Examined variables are shown as boxes and relationships as directional arrows with significant positive effects in blue, significant negative effects in red and non-significant paths in dotted grey based on a hypothesis driven SEM framework (Supplementary Figure 7). Standardized (significant) path-coefficients are shown next to each path with asterisks indicating significance (* P<0.05, ** P<0.01, *** P<0.001), path-width is scaled by coefficient size. Significant partial correlations are shown through grey, bi-directional arrows. The variation in species asynchrony and stability explained by fixed (left, marginal $R^2$) and fixed together with random model effects (right, conditional $R^2$) is shown in the corresponding boxes.

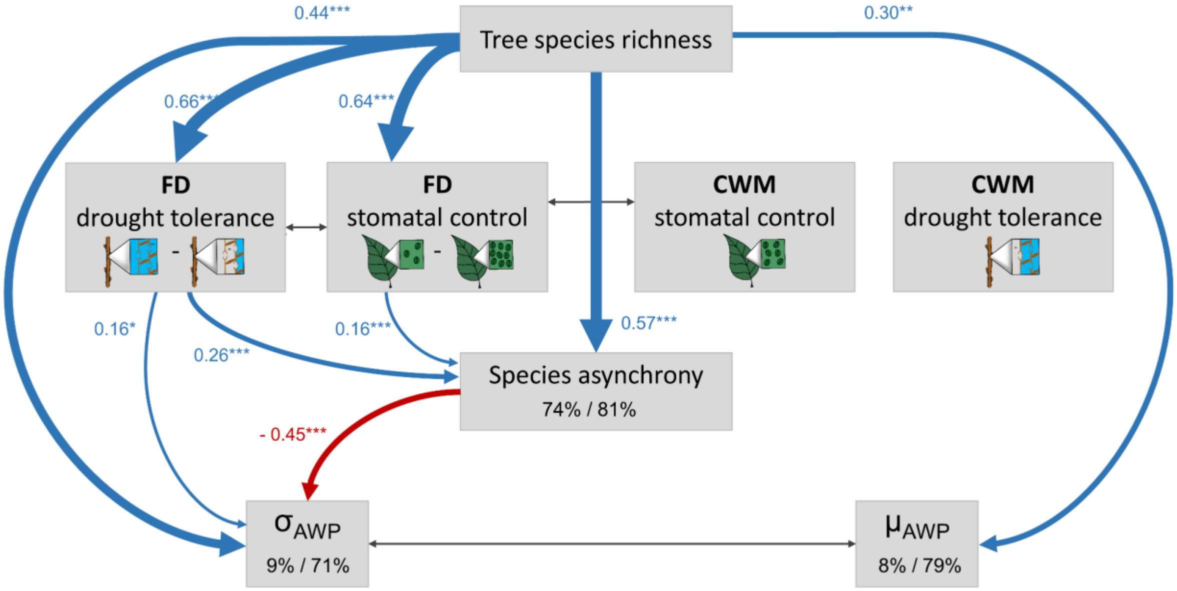


**Supplementary Figure 12** Partitioning of the direct and indirect effects of species richness, hydraulic diversity and community hydraulic means into overyielding and variance buffering effects of diversity. The structural equation model (SEM) is fit to data including monocultures and separates the hypothesized effects of diversity on stability (Fig. 3; Supplementary Figure 7) into effects on the temporal mean ($\mu_{AWP}$) and the temporal standard deviation of productivity ($\sigma_{AWP}$). Increases in $\mu_{AWP}$ enhance stability through overyielding – a higher productivity in mixtures vs monocultures - and decreases in $\sigma_{AWP}$ enhance stability through buffered variations in productivity. All drivers hypothesized to influence stability – species richness, functional diversity of stomatal control (FD stomatal control), functional diversity of drought tolerance (FD drought tolerance), the CWM of stomatal control (CWM stomatal control), the CWM of drought tolerance (CWM drought tolerance) and species asynchrony - were tested for their effects on $\mu_{AWP}$ and $\sigma_{AWP}$. Only significant pathways (P<0.05) are shown here to avoid overplotting (see Supplementary Figure 8 for the full model). The sketches schematically illustrate the trait gradients: water-spending vs water-saving stomatal control (few versus abundant stomata) and drought tolerance (high versus low cavitation resistance). The SEM fit the data well (Fisher's C=10.7, P=0.22, d.f.=8, n=375 plots). Data is based on a long, experimental species richness gradient ranging from monocultures to mixtures of 24 tree species. Examined variables are shown as boxes and relationships as directional arrows with significant positive effects in blue, significant negative effects in red and non-significant paths in dotted grey. Standardized (significant) path-coefficients are shown next to each path with asterisks indicating significance (* P<0.05, ** P<0.01, *** P<0.001), path-width is scaled by coefficient size. Significant partial correlations are shown through grey, bi-directional arrows. The variation in species asynchrony, $\mu_{AWP}$ and $\sigma_{AWP}$ explained by fixed (left, marginal $R^2$) and fixed together with random model effects (right, conditional $R^2$) is shown in the corresponding boxes.

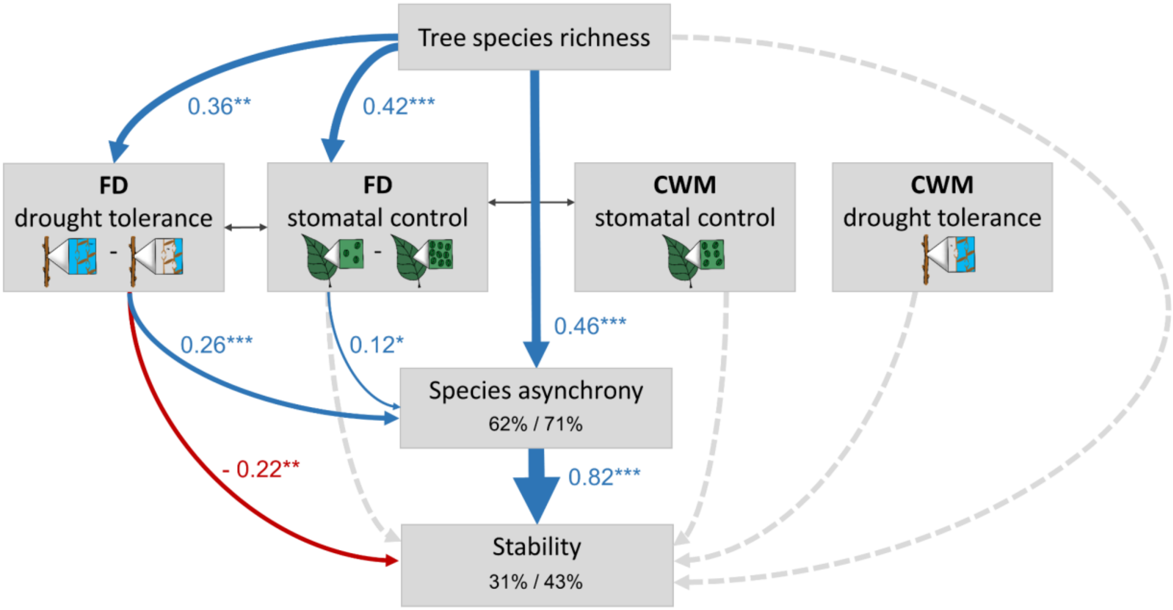


**Supplementary Figure 13** Direct and indirect effects of species richness, hydraulic diversity and community hydraulic means on stability. The structural equation model (SEM) is fit without square-root transformed species asynchrony and stability but with an exponential variance structure (varExp[10]) for log species richness and tests the effects of tree species richness, functional diversity of stomatal control (FD stomatal control) and functional diversity of drought tolerance (FD drought tolerance) on species asynchrony and stability. The sketches schematically illustrate the trait gradients: water-spending vs water-saving stomatal control (few versus abundant stomata) and drought tolerance (high versus low cavitation resistance). Functional diversity was calculated as abundance-weighted functional dispersion. Effects of community-weighted mean (CWM) traits are explored through testing the effect of the CWM of stomatal control (CWM stomatal control) and the CWM of drought tolerance (CWM drought tolerance) on stability. The SEM fit to the data was only marginally significant (Fisher's C=14.1, P=0.079, d.f.=8, n=218 plots). Data is based on a long, experimental species richness gradient with mixtures of 2, 4, 8, 16 and 24 tree species. Examined variables are shown as boxes and relationships as directional arrows with significant positive effects in blue, significant negative effects in red and non-significant paths in dotted grey based on a hypothesis driven SEM framework (Supplementary Figure 7). Standardized (significant) path-coefficients are shown next to each path with asterisks indicating significance (* P<0.05, ** P<0.01, *** P<0.001), path-width is scaled by coefficient size. Significant partial correlations are shown through grey, bi-directional arrows. The variation in species asynchrony and stability explained by fixed (left, marginal $R^2$) and fixed together with random model effects (right, conditional $R^2$) is shown in the corresponding boxes.

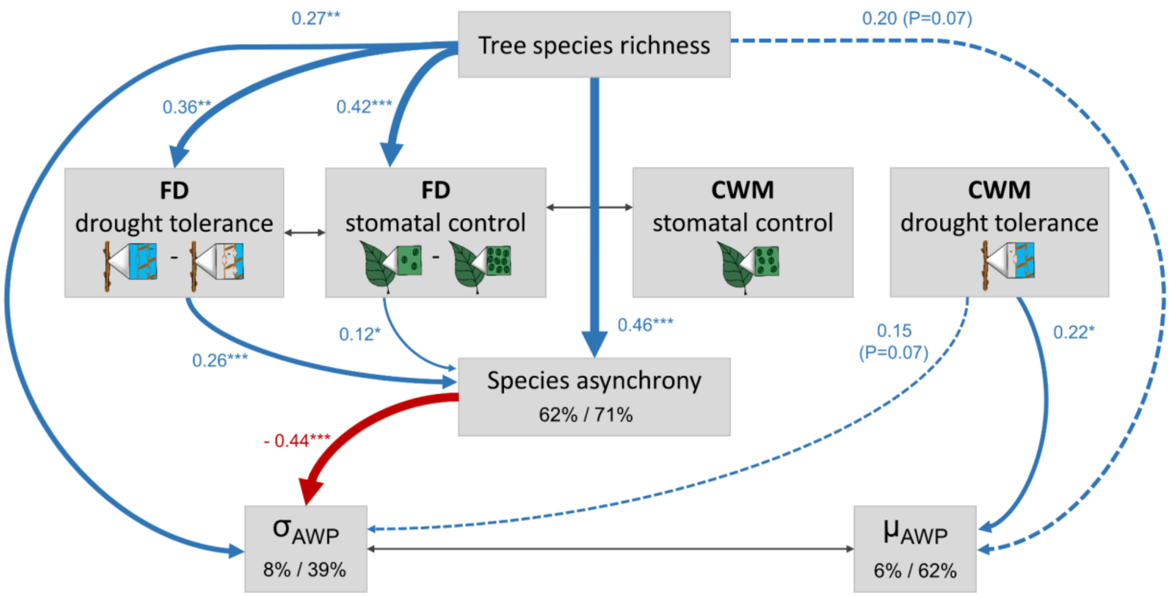


**Supplementary Figure 14** Partitioning of the direct and indirect effects of species richness, hydraulic diversity and community hydraulic means into overyielding and variance buffering effects of diversity. The structural equation model (SEM) is fit without square-root transformed species asynchrony and stability but with an exponential variance structure (varExp[10]) for log species richness and separates the hypothesized effects of diversity on stability (Fig. 3; Supplementary Figure 7) into effects on the temporal mean ($\mu_{AWP}$) and the temporal standard deviation of productivity ($\sigma_{AWP}$). Increases in $\mu_{AWP}$ enhance stability through overyielding – a higher productivity in mixtures vs monocultures - and decreases in $\sigma_{AWP}$ enhance stability through buffered variations in productivity. All drivers hypothesized to influence stability – species richness, functional diversity of stomatal control (FD stomatal control), functional diversity of drought tolerance (FD drought tolerance), the CWM of stomatal control (CWM stomatal control), the CWM of drought tolerance (CWM drought tolerance) and species asynchrony - were tested for their effects on $\mu_{AWP}$ and $\sigma_{AWP}$. Only significant pathways ($P<0.05$) are shown here to avoid overplotting (see Supplementary Figure 8 for the full model). The sketches schematically illustrate the trait gradients: water-spending vs water-saving stomatal control (few versus abundant stomata) and drought tolerance (high versus low cavitation resistance). The SEM fit to the data was only marginally significant (Fisher's C=14.1, P=0.079, d.f.=8, n=218 plots). Data is based on a long, experimental species richness gradient with mixtures of 2, 4, 8, 16 and 24 tree species. Examined variables are shown as boxes and relationships as directional arrows with significant positive effects in blue, significant negative effects in red and non-significant paths in dotted grey. Standardized (significant) path-coefficients are shown next to each path with asterisks indicating significance (* $P<0.05$, ** $P<0.01$, *** $P<0.001$), path-width is scaled by coefficient size. Significant partial correlations are shown through grey, bi-directional arrows. The variation in species asynchrony, $\mu_{AWP}$ and $\sigma_{AWP}$ explained by fixed (left, marginal $R^2$) and fixed together with random model effects (right, conditional $R^2$) is shown in the corresponding boxes.

### Supplementary Methods 1: Imputation of missing values

Missing diameter or height measurements in individual years are a common challenge in experiments utilizing inventory data sets with a high temporal resolution. Moreover, in 2016 a reduced sample of the central 6×6 trees was measured for all diversity levels except for the very intensively studied mixture plots (see ref[16] for details) in contrast to the central 12×12 trees in all other years. We therefore imputed missing tree diameter and height values of trees in a single year caused by forgotten measurements or the reduced sampling size in 2016 as detailed below, but only if the increment series was logical, i.e. $value_{x+1} \geq value_{x-1}$. To ensure that climate induced growth variability between years was preserved during imputation, we calculated site-specific (for site A and B) annual rates of individual tree growth that accounted for observed growth variability between years. We fitted a linear model that predicted annual increment in tree ground diameter (*gd*) by year and *gd* in the preceding year based on measured values of all trees that had complete and completely consistent increment series in all years of our observation period (2009-2019), that is, no missing and always positive increment values. We used the predicted annual increment to calculate rates of relative size change from one year to the next for all years as $r = \frac{i_x}{(i_x + i_{x+1})}$, where $i_x$ is the predicted *gd* increment in a year. These annual rates of change were than used to impute each missing measurement as $(v_{x+1} - v_{x-1}) * r_x + v_{x-1}$, where $v$ is the *gd* or *height* measurement in a year and *r* the rate of change. In total 2% of values (*gd*, *height* or both) were derived in this way. Hence, our imputation preserved observed annual tree growth changes as driven for example by climatic variability in between years while enhancing the completeness of our annual species and community level productivity estimates per plot.
